# Microplastics disrupt a cDC1-CD8/NKT regulatory circuit that restrains plasma cell accumulation after injury

**DOI:** 10.1101/2025.01.13.632166

**Authors:** Aditya Josyula, Daphna Fertil, Sonakshi Sharma, Isabella Horton, Devon R. Hartigan, Paige Rudy, Mayowa Amosu, Tran B. Ngo, Toluwaleke Fashina, Efua Maclean, Alessandra B. Coogan, Vanathi Sundaresan, Lucas dos Santos Dias, Michail S. Lionakis, Kaitlyn Sadtler

## Abstract

Traumatic injury exposes abundant self-antigens while simultaneously mounting a significant inflammatory response to counter infection and clear debris. While canonical regulatory T cells have been implicated in recovery from sterile injury, mechanisms of peripheral tolerance maintenance in clinically relevant models with wound contamination or material implants are unknown. Here, we identify a cDC1-dependent Ly49+ CD8/NKT-like immunoregulatory pathway that restrains autoreactive responses via Fas-mediated pruning of autoreactive immune cells. Microplastic contamination shifts this balance by limiting engagement of this pathway, thereby promoting the formation of tertiary lymphoid structures, local plasma cell generation, and autoantibody production, including insulin-reactive IgG and IgA. These findings identify a mechanism of immune regulation that restrains autoreactive responses after traumatic injury that is disrupted by persistent hydrophobic microplastic contaminants.

## INTRODUCTION

Traumatic tissue damage exposes self-antigens that are otherwise obscured from immune cells. Concomitantly, injury generates a robust inflammatory response to prevent infection and clear debris. Despite this potentially autoimmune microenvironment^1–3^, most patients recover without developing sustained autoreactive responses. Multiple layers of immune tolerance normally prevent this autoreactivity from initiating pathogenic immune responses. Although canonical FoxP3+ regulatory T cells are well established mediators of immune regulation in wound healing and regeneration in sterile models^4,5^, less is known about the acute recovery phase and clinically relevant injuries complicated by both therapeutically implanted materials and synthetic particulate debris from environmental contamination.

Traumatic injuries are frequently treated with a variety of degradable biomaterials to aid in wound healing and closure. Additionally, these traumatic wounds are exposed to synthetic polymers from implant wear particles and environmental contaminants. Microplastics are ubiquitous modern environmental contaminants that accumulate in human tissues^6,7^, though the extent of accumulation and its relationship with disease are still being investigated^8^. In contrast polymer wear particles generated from synthetic devices are established drivers of the foreign body reaction^9^. Whether these synthetic particulates alter the immunologic pathways that normally preserve self-tolerance in the context of persistent self-antigen exposure and wound inflammation are unknown.

Previously, using a volumetric muscle loss (VML) model in mice, we identified conventional dendritic cell type 1 (cDC1) which was recruited to injured skeletal muscle through XCL1 chemotaxis and was associated with T cell and macrophage immunoregulation with downstream effects on stem cell differentiation and tissue development^10^. These cells were enriched by a clinically used therapeutic biomaterial, decellularized extracellular matrix (ECM), associated with improved outcomes in wound healing and were inhibited by the presence of hydrophobic microplastics (polyethylene, PE). The foreign body response to PE results in robust neutrophil recruitment and persistent inflammation resulting in fibrosis^10,11^. Previous studies have also shown that naïve T cells continually traffic into chronically inflamed tissues and support tertiary lymphoid structures (TLS)^12,13^. Here, we show that traumatic injury generates autoreactive lymphocyte responses that are normally restrained by a cDC1-dependent Ly49+ CD8/NKT-like peripheral tolerance checkpoint. Microplastics shift this balance by reducing accumulation of cDC1s and CD8/NKT-like cells, allowing tertiary lymphoid responses and locally generated plasma cells to emerge. Taken together, these data highlight an immune checkpoint that limits acute autoreactive responses after injury and is bypassed by microplastic contaminants.

### CD8/NKT-like cells are depleted in wounds contaminated by microplastics

Biological scaffolds establish an immunoregulatory environment in damaged tissue after VML^10^. In contrast, fibrotic materials drive chronic inflammation and scarring^11,14^. Given the divergent phenotypes that result from materials in the wound environment, we investigated potential immunoregulatory phenotypes of muscle infiltrating lymphocytes at 7dpi using single cell sequencing (**Fig. 1a**). . We found that among the αβT cell populations was a group of cells with low *Cd8a* expression and high *Ikzf2* and *Klra* gene expression matching expression profiles of CD8 regulatory T cells^15–19^, MAIT cells and NKT cells^16,20^ (**Fig. 1a**). We confirmed the presence of CD8/NKT-like cells via flow cytometry where we detected HELIOS+Ly49+CD8+ (**Fig. 1b**) and CD8-(**Fig. 1c**) T cells using an Ly49 inhibitory receptor antibody cocktail including Ly49A/C/I/F/H. These cells represented a major fraction (∼25%) of all αβT cells and expressed transcripts encoding the cDC1 chemoattractant *Xcl1*. Notably, the recruitment of these cells was enriched upon ECM_tx_ and attenuated by PE_tx_ after VML.

**Figure 1.**
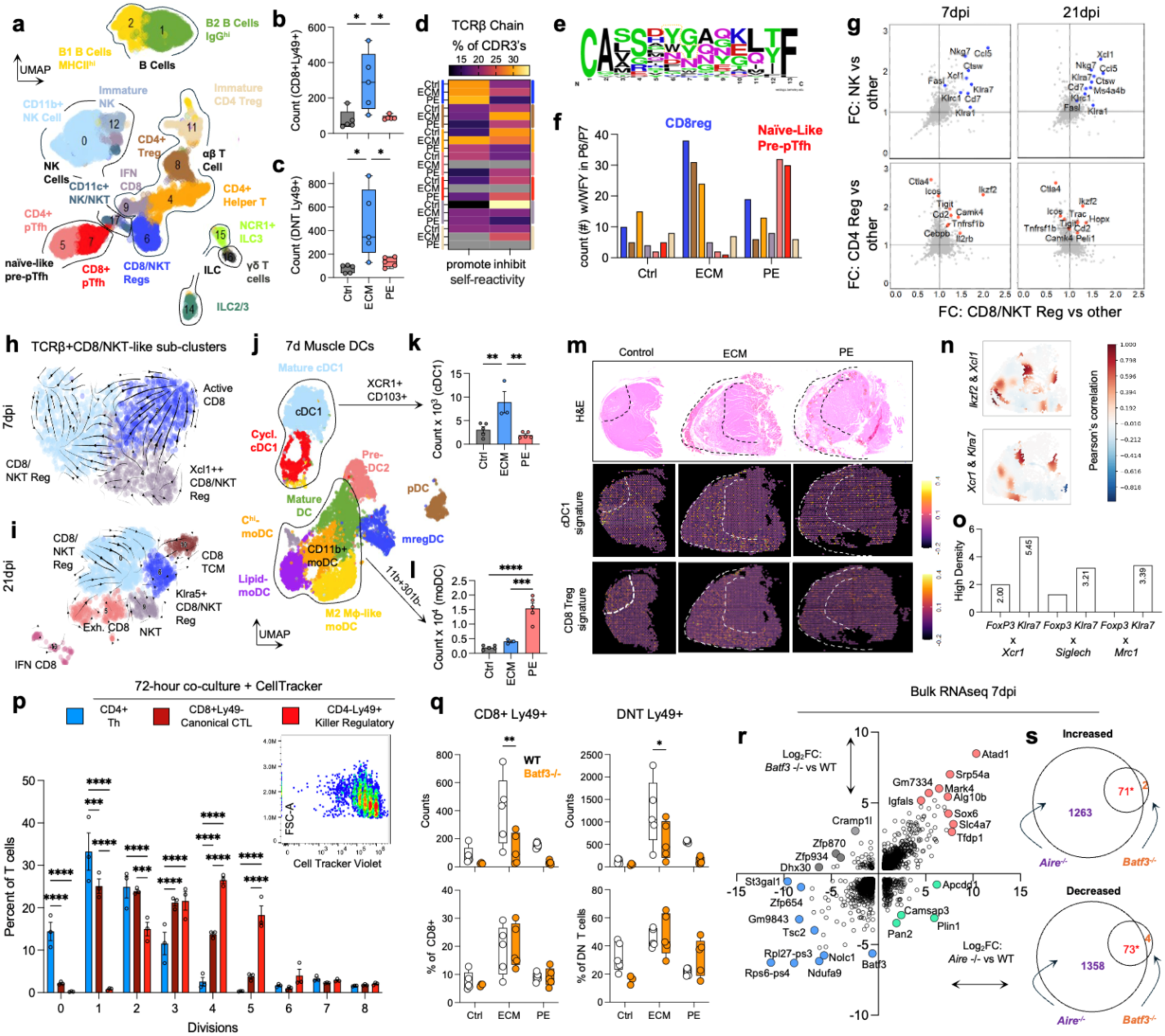
| cDC1s support activation of CD8/NKT-like cells and are depleted in wounds contaminated by microplastics. (**a**) UMAP display of Seurat clusters of CD11blo immune cells in muscle injury at 7 days post-injury. (**b-c**) Flow cytometry quantification of TCRβ+ CD4-(**b**) CD8a+ Ly49(A/C/I/F/H1)+ or (**c**) CD8a-Ly49(A/C/I/F/H1)+ in muscle wound at 21 days post-injury. (**d**) CDR3 self-reactivity index by hydrophobic residues in P6/P7 of TCRβ 13mer sequences of different Seurat clusters at 7 days post injury. (**e**) Logo plot of TCRβ 13mer sequences of Ly49+ CD8/NKT-like population at 7 days post-injury (**f**) Count of aromatic (tyrosine, phenylalanine, tryptophan) residues at P6/P7 in different T cell clusters at 7 days post-injury. (**g**) Transcriptional similarities between Ly49+ CD8/NKT-like population and classic NK cells (top, blue) and classic CD4+ Tregs (bottom, red) at 7 days post-injury. (**h**-**i**) RNAvelocity of CD8/NKT-like seurat clusters at 7 (**h**) and 21 (**i**) days post-injury. (**j**) UMAP display of Seurat clusters of MHCII+CD11bloCD11c+ dendritic cells at 7 days post-injury. (**k-l**) flow cytometry quantification of (**k**) CD103+XCR1+ cDC1s and (**l**) CD11b+CD301b-CD206-SiglecH-moDCs. (**m**) spatial transcriptomics of cDC1 and Ly49+ CD8/NKT-like signature (**n**) colocalization of Ly49+ CD8/NKT-like gene *Ikzf2* with cDC1 chemotactic signal *Xcl1* and cDC1 marker *Xcr1* with Ly49+ CD8/NKT-like gene *Klra7*. (**o**) Quantification of colocalization by high pixel density area of CD4 T regulatory (*Foxp3*) and Ly49+ CD8/NKT-like (*Klra7*) with cDC1 (*Xcr1*), pDC (*Siglech*), and M2 macrophage (*Mrc1*) representative genes. (**p**) T cell division measured by CellTracker Violet after co-culture of bone marrow derived cDC1s with CD4+ T cells, canonical CD8+ T cells, or CD4-Ly49+ CD8/NKT-like for 72 hours. (**q**) CD8+ and CD8-CD8/NKT-like populations in WT and cDC1-deficient (*Batf3^-/-^*) mice. (**r**) Double volcano plot of bulk RNAseq of muscle tissue at 7 days post injury in *Batf3^-/-^* and *Aire^-/-^* mice compared to WT mice. (**s**) Overlap of significantly increased (top) or decreased (bottom) genes in *Aire^-/-^* and *Batf3^-/-^* versus WT mice. Data are range with IQR (b,c,k,l,p,q) or mean and standard error (o). ANOVA with Tukey post-hoc correction for multiple comparisons, **** = p < 0.0001, *** = p < 0.001, ** = p < 0.01, * = p < 0.05, n = 3 – 5 mice representative of at least two independent experiments (flow cytometry and cell culture), pooled n = 10 (lymphocytes) and n = 5 (dendritic cells) for single cell sequencing, and n = 3 mice for bulk RNA sequencing.

A detailed evaluation of CDR3 sequences showed differences in amino acid enrichment at different positions within the CDR3 (**Fig.1e**). When evaluating 13-mer CDR3s (TCRβ chain) we found an overall CA(AA3-12)F pattern in the majority of CDR3s evaluated with asparagine, glycine, and serine being enriched within the central amino acids. While amino acids on the periphery were more likely to be hydrophobic (positions 2, 3, 13) central amino acids tended towards polar uncharged classifications. Given these variations, we evaluated potential patterns of amino acids that were previously associated with promoting and inhibiting self-reactivity^21^. We generated self-reactivity indices for all clusters based on their amino acids at positions 6 and 7 of the CDR3 as reported by Stadinski et al^21^. We found that over 25% of clones detected within the Ly49+ CD8/NKT-like cluster were identified as “self-reactive” as opposed to other groups with lower indices (**Fig.1d**). Overall, these predicted self-reactive clones showed an enrichment of aromatic amino acids like tyrosine and tryptophan in positions 6 and 7, which was the highest in ECM_tx_ in comparison to control and PE_tx_ injuries (**Fig.1f**). Additionally, we computed a cysteine index (**SFig. 1a**) at the CDR3 apex for all tissue infiltrating T cells at 7 and 21 dpi. Ly49 expressing cells in tissue had significant enrichment of the cysteine index further supporting potential self-reactivity. Notably, these cells were not restricted in TRA or TRB gene usage and displayed a wide range of TRA and TRB gene usage (**SFig. 1b**).

To further understand their phenotype, we compared CD4+ FoxP3+ conventional T regulatory cells and CD8/NKT-like cells, we noted a shared phenotype comprising *Il2rb, Cd27, Tigit*, and *Ikzf2.* A similar analysis with NK cells showed a shared expression pattern comprising, *Ccl5, Nkg7, Xcl1* and *Klra7* (**Fig.1g**). To assess any possible transcriptional dynamics among various autoreactive CD8/NKT-like T cell subsets within tissue, we subclustered all activated CD8 αβT cells and NKT cells and performed RNA velocity. At early activation stages (7 dpi) we observed a strong transcriptional trajectory from *Xcl1* expressing CD8 T cells branching into *Ikzf2*-CD8/NKT-like cells as well as *Ikzf2*-canonical CD8 T cells (**Fig.1h**). No strong trajectories were observed at 21 dpi (**Fig.1i**). Taken together, our data suggests that self-reactive Ly49+ CD8/NKT cells that share transcriptional programs with NK cells and Foxp3+ regulatory T cells are recruited to damaged tissue and attenuated by microplastics (PE_tx_).

### cDC1s support activation of CD8/NKT-like cells

We confirmed our previous findings of cDC1 accumulation in damaged tissue and treatment dependent enrichment using flow cytometry, spatial transcriptomics and single cell sequencing (**Fig.1j-k** and **SFig. 2**). Additionally, we found that PE_tx_ enriched clusters resembling monocytic DCs specifically those that express *Slc7a11*, *Fas, Ccr7* and *Clec4d* in tissue (**Fig.1l** and **SFig. 2a-c**). In contrast, pre-cDC2s and scavenger receptor high DCs were enriched in ECM and depleted in PE (**SFig. 2d-e**).

**Figure 2.**
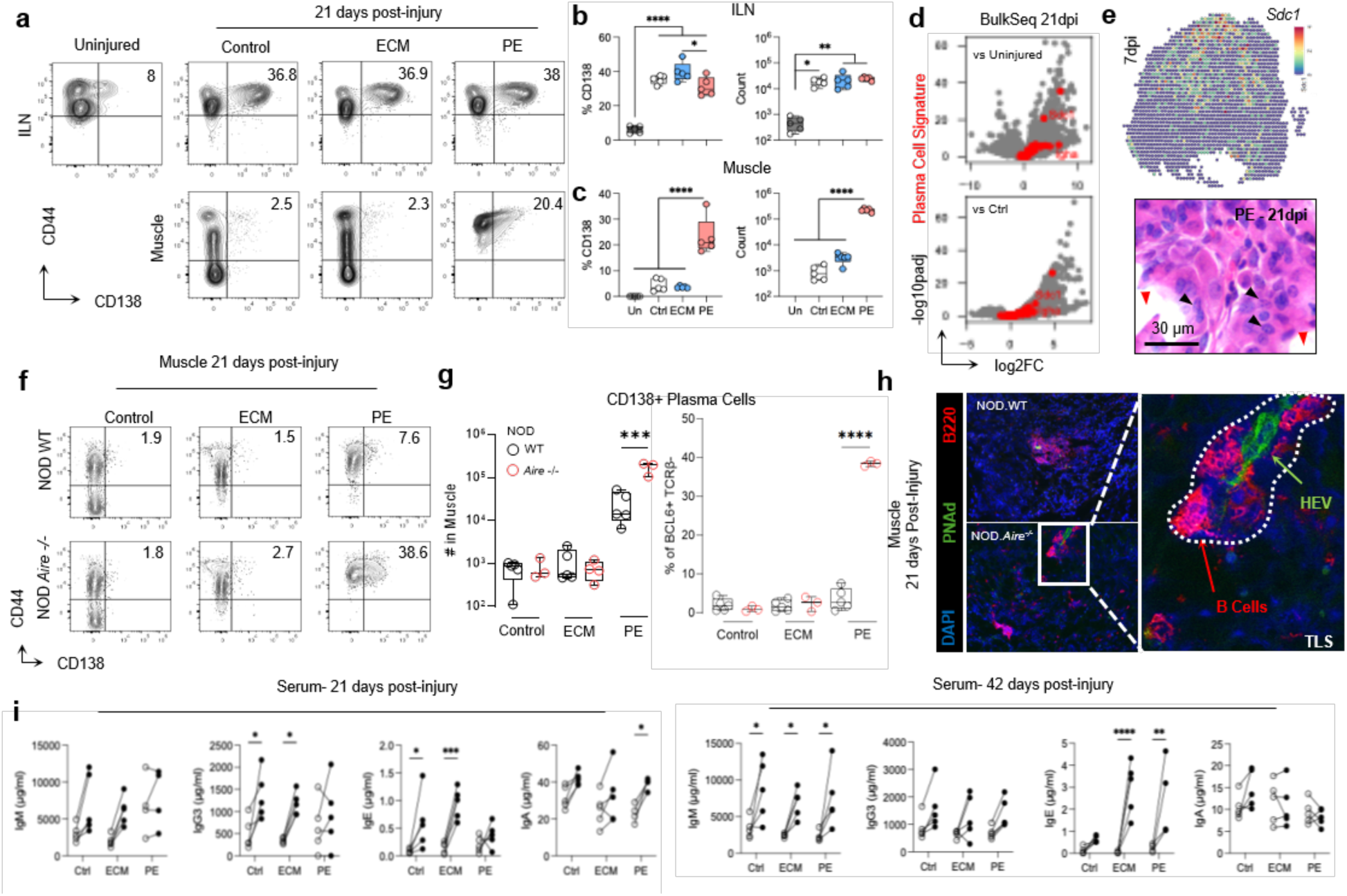
| Microplastics induce local plasma cell generation in damaged muscle and trigger autoimmunity. (**a**) Representative flow cytometry plots of plasma cells in inguinal lymph node (ILN) and muscle tissue at 21 days post-injury with control injury, extracellular matrix (ECM) treatment and polyethylene (PE) microplastic treatment. (**b-c**) Quantification of CD138+BCL6+ plasma cells at 21 days post-injury in (**b**) inguinal lymph node and (**c**) injured muscle tissue. (**d**) Bulk RNA sequencing of muscle tissue at 21 days post-injury of PE-treated muscle. Plasma cell signature genes highlighted in red. (**e**) Spatial RNA sequencing of *Sdc1* (encoding CD138) and histological identification (black arrows) at 7 days post-injury in PE-treated (red arrows) muscle. (**f**) Representative flow cytometry plots of plasma cells in NOD and NOD-*Aire^-/-^* mice at 21 days post-injury. (**g**) Quantification of data shown in panel f. (**h**) Immunofluorescence staining of PNAd and B220 for tertiary lymphoid structure (TLS) identification in NOD and NOD-*Aire^-/-^*mice. (**i**) Antibody concentrations in serum at 21- and 42-days post-injury. Statistical significance is by ANOVA with Tukey post-hoc correction for multiple comparisons, **** = p < 0.0001, *** = p < 0.001, ** = p < 0.01, * = p < 0.05, n = 3 – 5 mice in b,c and g. Comparisons indicated in I are by paired t-test.

Given that CD8/NKT-like cells showed significant *Xcl1* expression, and that cDC1s can cross-present antigen to CD8+ T cells, we hypothesized that cDC1s communicate with these cells in tissue. To test this, we first developed a transcriptional signature comprising *Xcr1*, *Itgae* and *Itgax* and mapped it onto spatial transcriptomics spots of injured muscle and indicated treatments (**Fig.1m**). We also performed a similar analysis for CD8/NKT-like cells and mapped a signature of *Il2rb*, *Xcl1*, *Ccl5, Gramd3, Klrb1c* and *Klra1/4/7* genes onto spatial transcriptomic spots (**Fig.1m and SFig. 2f**). We noticed accumulation of both signatures within areas of tissue damage particularly in ECM_tx_ where both populations were enriched. Further, we parsed the signature markers *Xcr1*/ *Klra7* and *Ikzf2/Xcl1* and assessed spatial colocalization (**Fig.1n**). We observed multiple hot-spots of colocalization within tissue suggesting cell-cell contact which we confirmed was not solely attributable to abundance by mapping colocalization of negative control markers such as the macrophage receptor *Mrc1* as well as plasmacytoid DC marker *Siglech* which showed lower colocalization (**Fig.1o**).

Having observed co-localization, we next asked if the interaction between cDC1s and CD8/NKT-like cells gave rise to Ly49+ CD8/NKT-like cell expansion which is a characteristic response to antigen presentation events *in vitro*. To test this, we co-cultured bone marrow derived, Flt3-L induced cDC1s pulsed with ECM proteins and CD8/NKT-like cells sorted from inguinal lymph nodes and spleens of mice with ECM_tx_ injuries. Cell trace violet dilutions of CD4+ T cells, CD8+Ly49-T cells and CD8/NKT-like cells showed selective proliferation of CD8/NKT-like cells 72 hours after co-culture (median = 2 divisions vs 4 divisions, p<0.0001) (**Fig.1p**). Next, to identify potential pathways of cDC1-CD8/NKT-like cell crosstalk, we extracted highly expressed markers and used those and inputs for molecular pathway analysis using the STRING database. We found enrichment of IL-18 signaling, T cell activation, and chemokine-receptor interactions. Overall, through STRING and CellChat analyses, we identified 9 distinct communication networks between cDC1s and CD8/NKT-like cells which could result in priming and downstream effector functions (**SFig. 3a and b**). Lastly, to test the effect of cDC1 depletion on CD8/NKT-like cells, and to investigate downstream events in mice lacking the cDC1-CD8/NKT-like immune checkpoint, we performed VML with all treatment groups in *Batf3^-/-^* mice which we previously confirmed to lack cDC1s^10^. These mice showed a general decrease in Ly49+ CD8/NKT-like populations in the muscle after VML with a significant decrease in mice which received ECM_tx_ which further supports the hypothesis of cDC1 communication with CD8/NKT-like cells (**Fig.1q**). To extend the relevance of our findings, we analyzed publicly available single cell RNA sequencing datasets to look for signature markers of Ly49+ CD8/NKT-like cells (**SFig. 3c**) in other mouse models. Indeed, Ly49+ CD8/NKT-like cells represented ∼10% of αβT cells in the context of skin injury and ∼25% in the context of inflammation induced damage of mouse myometrium. These findings suggest that Ly49+ CD8/NKT-like cell accumulation in the site of injury is conserved across tissues (**SFig. 3d**). Additionally, we also found that an analogous (KIR+) cell type comprises approximately 5% of αβT cells in human skeletal muscle (**SFig. 3e**).

**Figure 3.**
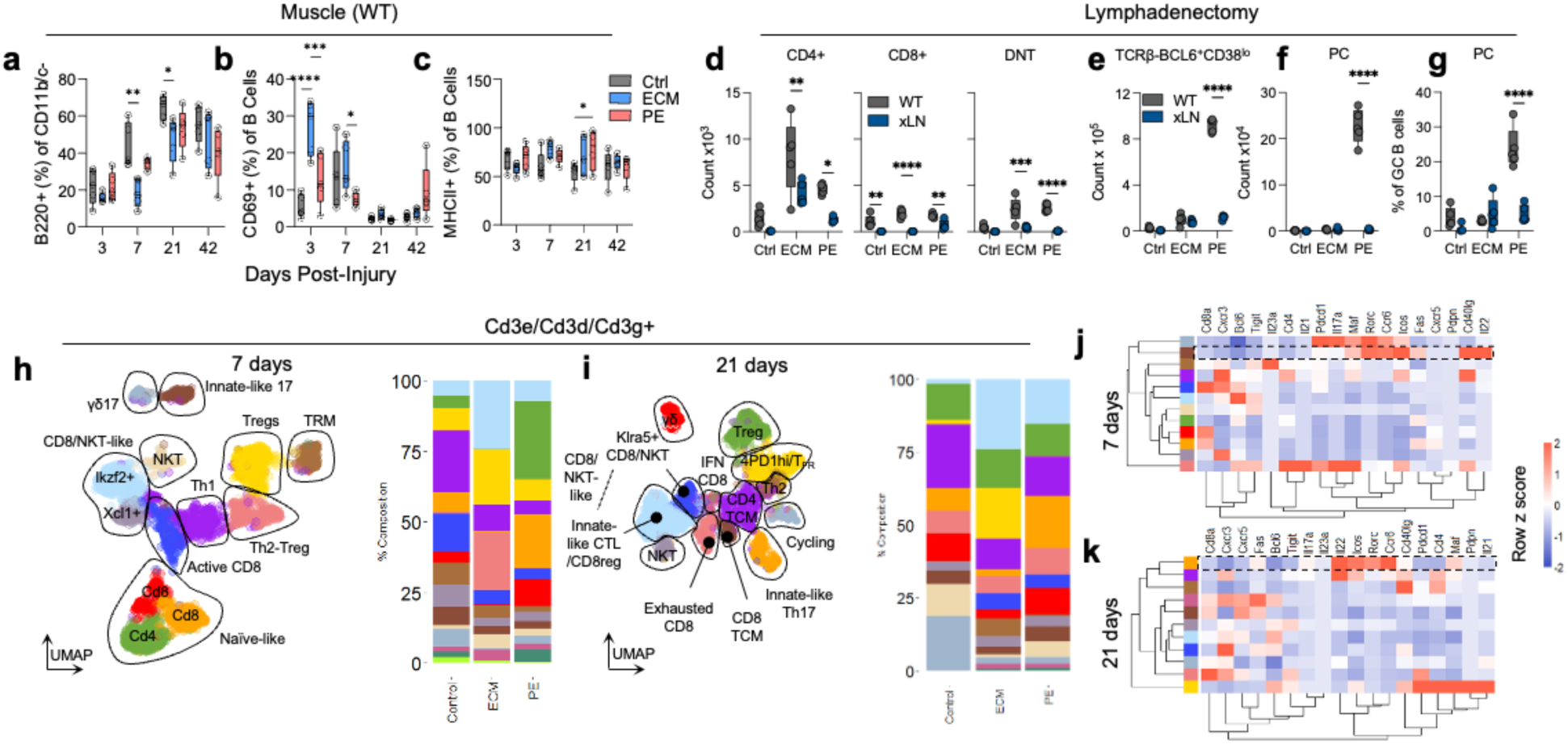
| T cells from draining lymph node of damaged tissue drive local plasma cell differentiation. (**a-c**) B220+ B cell frequency within the local muscle tissue as a percent of CD11b/CD11c-cells (excludes plasma cells) (**b**) early active CD69+ as a percent of B220+ B cells (**c**) MHCII+ as a percent of B220+ B cells. (**d**) αβT cell recruitment to injured muscle at 21 dpi in mice with intact lymph nodes and mice which received lymphadenectomy prior to VML. (**e-g**) Plasma cell abundance and fractions after lymphadenectomy prior to muscle injury surgery. (**e**) Counts of germinal center B cells (**f**) Counts of plasma cells (**g**) Percent of Germinal center B cells converted to plasma cells. Grey = lymph node intact, Blue = lymphadenectomy “xLN”. (**h-i**) UMAP display of seurat clusters from single cell RNA sequencing of T cells (Cd3e/Cd3d/Cd3g+) in the injured muscle tissue at (**h**) 7 days and (**i**) 21 days post-injury. (**j-k**) Type-17 self-reactive cell gene signature in Seurat clusters at (**j**) 7 and (**k**) 21 days post-injury.

To understand the contribution of cDC1-dependent immune responses in restraining the autoimmunity, we performed bulk RNA sequencing of injured muscle tissue from *Aire^-/-^*, *Batf3^-/-^* (cDC1-deficient) and WT mice at 7dpi (**SFig. 4a**). Comparing differentially expressed genes in *Aire^-/-^* and *Batf3^-/-^* to WT controls revealed a highly overlapping expression pattern including upregulated *Atad1*, *Srp54a, Mark4* and *Slc4a7* and downregulated *Tsc2, Ndufa9*, and *Nolc1* (**Fig.1r**). Broadly, tissue level differential expression of genes in *Batf3^-/-^* vs WT were nearly fully recapitulated within *Aire^-/-^* phenotype (71 of 73 genes increased, 73 of 77 genes decreased) (**Fig.1s**). Indeed, in *Aire^-/^*^-^ mice, there was elevated immune cell infiltration of injured tissue (**SFig. 4b-c**) and accompanying CD62L shedding in T and B cells suggesting broad lymphocyte activation (**SFig. 4d**). These data suggest that mice that lack central tolerance (*Aire^-/-^*) or cDC1 dependent peripheral regulation (*Batf3^-/-^*) converge on mechanisms that promote T cell infiltration and inflammation after tissue damage. Further, the cDC1 dependent peripheral regulation in damaged tissue is significantly attenuated upon exposure to microplastics.

**Figure 4.**
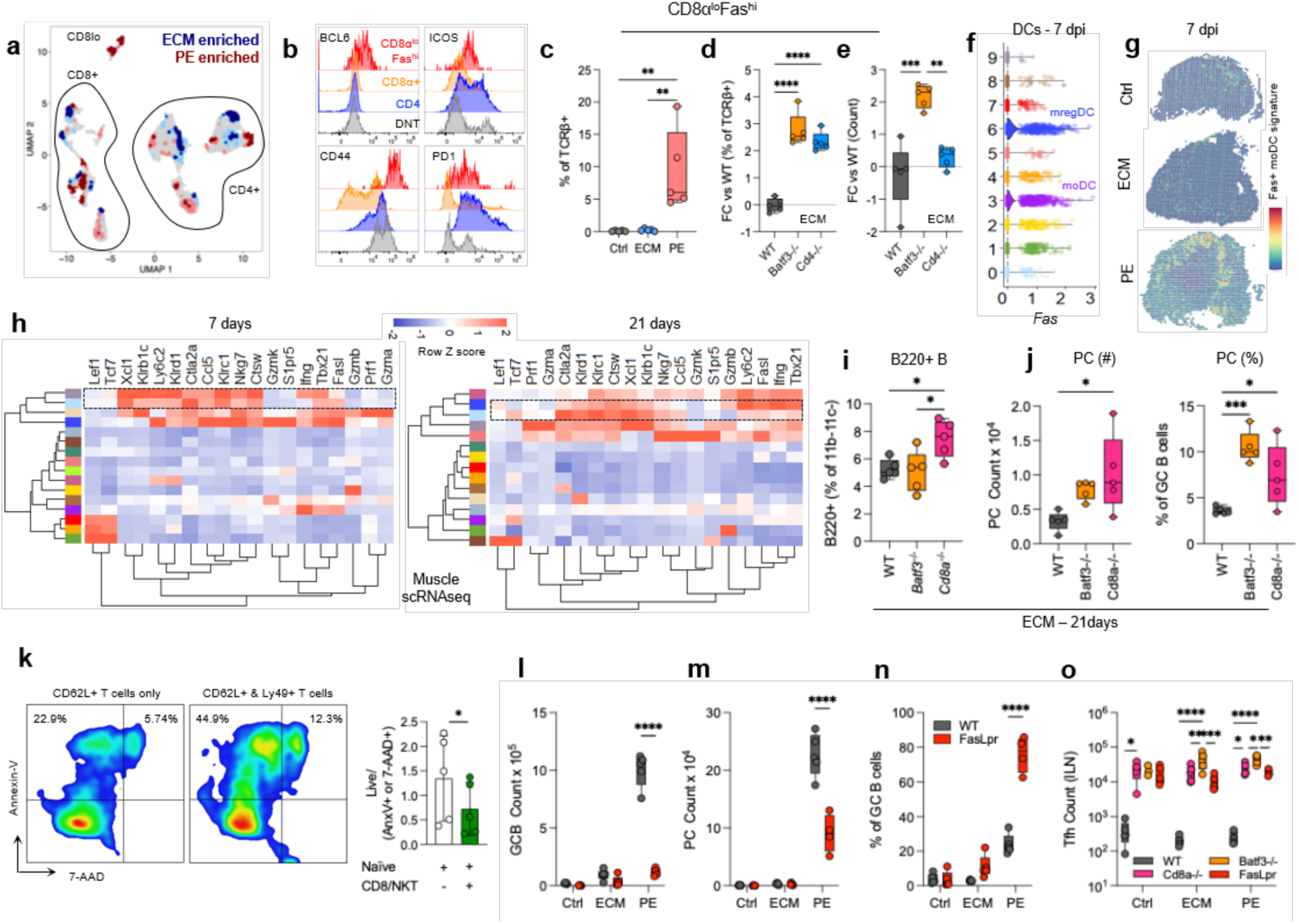
| cDC1 and Ly49+ CD8/NKT-like cells contribute to Fas mediated restriction of T follicular helper cells in lymph nodes and plasma cell conversion in TLS after tissue damage. **(a)** TREX analysis of T cell flow cytometry characterization of ECM_tx_ versus PE_tx_ injuries. **(b**) CD8a^lo^Fas^hi^ population identified as enriched in PE_tx_ mice. (**c**) Quantification of CD8a^lo^Fas^hi^ cells. (**d-e**) Presence of CD8a^lo^Fas^hi^ cells in ECM treated mice in WT (grey) *Batf3^-/-^* (orange) and *Cd4^-/-^* (blue) mice as a (d) percent of TCRβ+ cells or (e) total count. (f) Violin plot of *Fas* expression in DC Seurat clusters, cluster 3 = moDCs with lipid-like signature (g) Spatial RNA sequencing of moDC gene signature at 7 days post-injury. (h) Heatmap of z-scores of killer-like gene expression in Seurat clusters from scRNAseq at 7- and 21-days post-injury. (i-j) B cells in ECM_tx_ WT, *Batf3^-/-^* and *Cd8a^-/-^* mice (i) B220+ B cells. (j) BCL6+CD138+ plasma cells and BCL6+CD138-germinal center B cells (GCB). (k) Cell death of CD62L+ T cells (sorted from PEtx muscle) after co-culture with Ly49 inhibitory receptor positive CD8/NKT-like cells (sorted from ECM treated muscle) by staining with Annexin V and 7-AAD to detect apoptotic and dead cells. (l-n) GCB and plasma cell infiltration into the muscle tissue of WT (grey) and Faslpr (red) mice. (l) GCB count (m) plasma cell count (**n**) plasma cells as a percent of total germinal center B cells. (**o**) CD4+ Foxp3-CXCR5+ PD-1+ T follicular helper cells in the ILN of WT (grey), Batf3-/- (orange), Cd8a-/- (pink) and Faslpr (red) mice.

Furthermore, we found that CD8/NKT-like cells (**SFig. 5a**) were present within the peripheral blood, spleen and inguinal lymph nodes at varying frequencies (**SFig. 5b**). Interestingly, HELIOS expression within these cells, at the protein level, was drastically reduced upon activation across WT, *Batf3-/-* and *Cd8a-/-* mice (**SFig. 5c**). Unlike CD4+ FoxP3+ Tregs, these cells did not gain BCL6 expression after activation (**SFig. 5d**). Further, in mice that lack *Batf3* or *Cd8a* we observed an increase in CD4+ Foxp3+ Tregs (**SFig. 5e**) in the damaged tissue at 7dpi indicating a possible compensatory mechanism.

**Figure 5.**
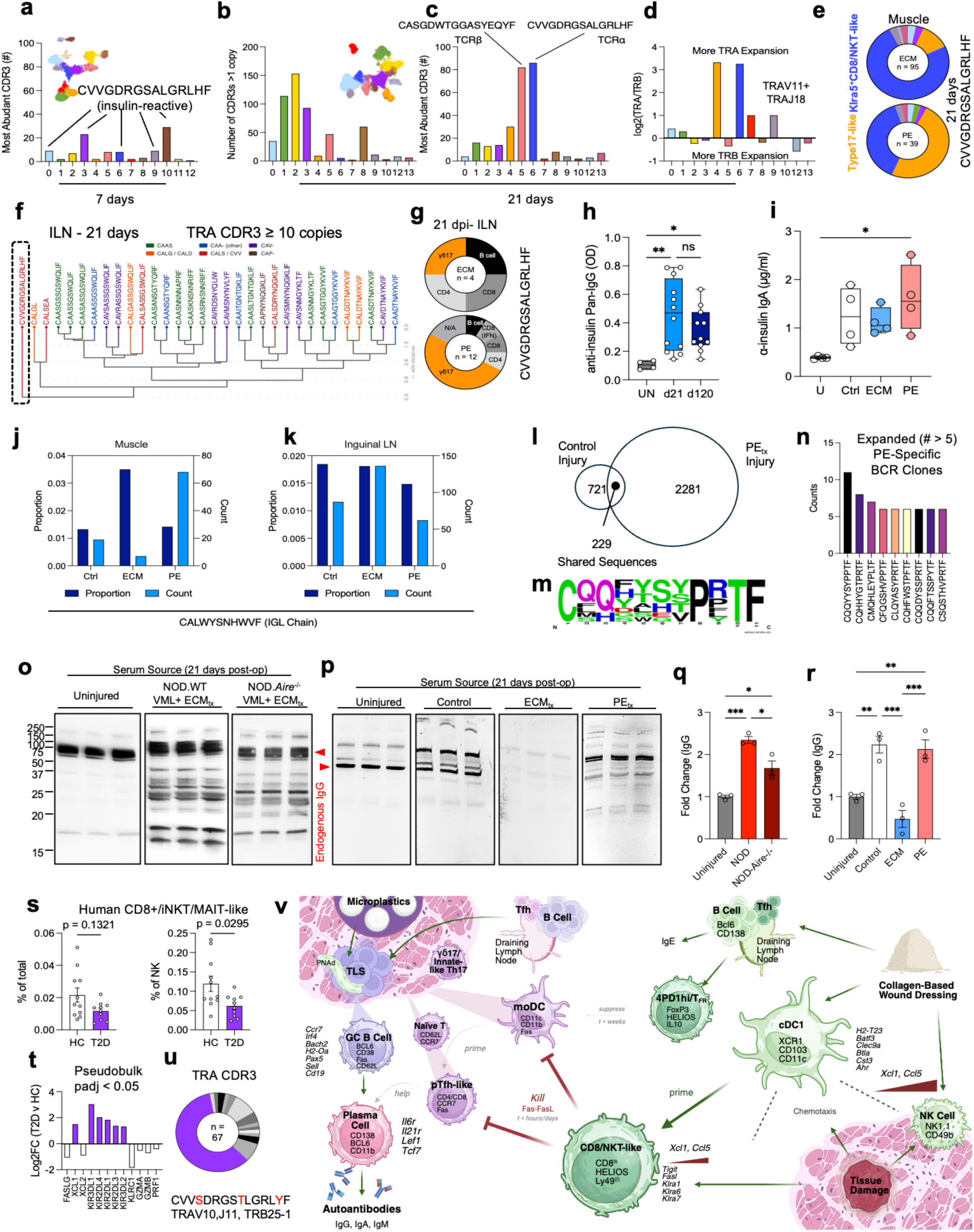
| Skeletal muscle damage induces self-antigen specific T and B cell responses locally. (**a**) CDR3 (TRA or TRB) expansion in muscle-infiltrating T cells at 7 days post-injury displayed as count of highest expanded CDR3 per cluster (**b**) Overall oligoclonal expansion as number of CDR3s with more than 1 copy detected. (**c**) Count of the most abundant CDR3 sequence. (**d**) Relative expansion of TRA or TRB by cluster as a ratio of TRA highest expanded clone count over TRB highest expanded clone count. (**e**) Seurat cluster presence of CVVGDRGSALGRLHF CDR3 sequence in T cell clusters. (**f**) Highly expanded (≥10 copies) CDR3 TRA sequences in inguinal lymph node plotted by edit distance. (**g**) Seurat cluster presence of CVVGDRGSALGRLHF CDR3 sequence in overall ILN clusters. (**h**) Serum levels of anti-insulin IgG and (**i**) IgA in indicated treatments and timepoints. (**j-k**) Enrichment of highest enriched CDR3 sequence in (**j**) muscle and (**k**) inguinal lymph node. (**l**) Overlap of CDR3 sequences between control injury and PE_tx_ injury. (**m**) CDR3 logo plot of shared sequences with control injury. (**n**) Expanded PE-specific BCR CDR3 sequences. (**o-p**) IgG antibodies detected in serum of (**o**) NOD and (**p**) WT C57BL/6 mice. (**q-r**) Quantification of (**q**) NOD and (**r**) WT C57BL/6 blots. (**s**) Prevalence of CD8+NKT-like cells in peripheral blood of human type 2 diabetes patients (T2D, purple) versus healthy controls (HC, white) (**t**) Pseudobulk comparison of CD8+/NKT-like cells in HC vs T2D (**u**) Prevalence of CVVSDRGSTLGRLYF sequence in PBMC clusters. (**v**) Schematic of tolerogenic cascade mediated by Ly49+ CD8/NKT-like cells after traumatic injury. Cell culture n = 3 representative of 3 independent experiments. Flow cytometry n = 4 – 5 representative of at least 2 independent experiments. Antibody titers n = 3 – 5 representative of at least 2 independent experiments. ANOVA with Tukey post-hoc correction for multiple comparisons, **** = p < 0.0001, *** = p < 0.001, ** = p < 0.01, * = p < 0.05. Panel (**v**) was made using BioRender.

### Microplastics induce local plasma cell generation in damaged muscle

Previous reports of Ly49+ HELIOS+ CD8+ T cells in mice and humans suggest that a function of these cells is to regulate pathogenic T cells to prevent autoimmunity^19,22^. Since, PE_tx_ attenuated these CD8/NKT-like cells (**Fig.1a-b**) that closely resemble this previously reported phenotype, we investigated T and B cell populations that are differentially recruited to PE treated mice after tissue damage which could potentially lead to autoimmunity.

While skeletal muscle tissue does not harbor a significant population of lymphocytes under homeostasis, it is known that various T and B cell subsets are recruited upon tissue damage alongside antigen presenting cells^11,23^. Using a flow cytometry panel, we tracked the accumulation and activation status of T and B cells in the muscle and draining lymph node over a period of 6 weeks (**SFig. 7a**) and observed continual recruitment of naïve-like T cells into damaged muscle. Therefore, we hypothesized that lymphocytes may organize into tertiary lymphoid structures (TLS) to maintain a local lymphocyte population. A hallmark of TLS is the presence of germinal center-like B cells (B220^lo^CD38^lo^CXCR5^+^Fas^+^BCL6^+^)^24,25^. We observed that while tissue damage across treatment groups led to GC-like B cells in tissue, mice which received PE had a classical phenotype of plasma cells (CD138^+^CD44^+^), arising from GC-like B cells (**Fig.2 a-c**). Additionally, VML induced significant plasma cell generation in draining lymph nodes as compared to uninjured mice but not in the spleen (**Fig.2 a-b** and **SFig. 6a-b**). By bulk RNA sequencing of muscle tissue (PE vs control and PE vs uninjured; 21 dpi) we confirmed that plasma cells differentiate locally in damaged tissue with PE (**SFig. 6c-f**). GSEA analysis comparing mice treated with PE to control injury and ECM treatment (ECM_tx_) showed two independent plasma cell marker gene sets enriched with PE (**SFig. 6g-h**). Furthermore, a plasma cell gene signature comprising heavy chain genes, *Sdc1, Prdm1* and *Irf4* was enriched in damaged muscle with PE (**Fig.2d**). *Sdc1* expression within spatial transcriptomic spots from histological sections of mice with PE in wounds showed elevated and material specific spatial distribution of plasma cells at 7dpi (**Fig.2e** and **SFig. 6i-j**). Histologically, we identified plasma cells within injured muscle surrounding PE at 21dpi (**Fig.2e**). Together, these data revealed significantly elevated plasma cell enrichment with PE in injured muscle tissue.

### Microplastics synergize with *Aire* deletion to potentiate local plasma cell generation

To extend the relevance of these observations, we investigated ectopic plasma cell generation in non-obese diabetic (NOD) WT and NOD.*Aire^-/-^* mice which closely model pathological autoimmunity in humans^26,27^. While PE treatment modestly increased local plasma cell numbers in NOD WT mice, we observed marked synergy in plasma cell generation in PE exposed NOD.*Aire^-/-^* mice (**Fig.2 f-g**). Indeed, in both NOD WT and *Aire^-/-^* mice, we observed B cell clusters surrounding high endothelial venules (HEV), which are definitive indicators of TLS (**Fig.2h**) but only PE exposure led to plasma cell generation in injured muscle. Additionally, in WT C57BL/6 markers that support the formation and maintenance of TLS^28,29^ (*Cd300lg*, *Lta* and *Ltb*) were present at the site of injury at 7 dpi (**SFig. 6k**). Like C57BL/6 WT mice, both NOD WT and NOD *Aire^-/-^* mice which underwent VML generated plasma cells in inguinal lymph nodes (**SFig. 6l**).

Given that we observed plasma cell generation both within damaged tissue and draining lymph nodes, we investigated systemic differences in antibody titers at 21dpi and 42dpi in WT C57BL/6 mice after VML across treatments as compared to pre-injury levels (**Fig.2i**). In agreement with previous reports^30,31^, injury alone elevated circulating IgE and IgM. In contrast to control injury and ECM_tx_, IgA was significantly elevated in mice with PE treatment (**Fig.2i**). Somatic hypermutation within multiple isotypes was evident in draining lymph nodes with the exception of IgE producing plasma cells which were not detectable lymph nodes via sequencing (**SFig. 6m**). Notably, plasma cells generated locally in PE mice co-expressed CD138 and CD11b which is widely accepted to indicate potent albeit transient IgA secretion by plasma cells^32^.

### T cells recruited from draining lymph nodes are required for plasma cell differentiation in damaged tissue

Since microplastics promoted robust local plasma cell differentiation and CD8^+^Ly49^+^HELIOS^+^ T cells have been shown to prune helper T cells^33^, we sought to identify helper T cell subsets in tissue that enabled plasma cell differentiation. We observed early B cell activation at 3-7 dpi and retention of MHC-II+ B cells in tissue alongside a naïve T cell accumulation in the muscle for up to 42 days after VML (**Fig.3 a-c** and **SFig. 7a**). Skeletal muscle tissue does not harbor a local pool of naïve T cells or helper cells and therefore we hypothesized that T cells are recruited from the blood, draining lymph node or distal secondary lymphoid organs to facilitate plasma cell differentiation. To investigate this, we first probed T cells, GC-like B cells and plasma cells within damaged tissue of mice that underwent lymphadenectomy of quadricep muscle-draining inguinal lymph nodes prior to VML (xLN) as compared to those from damaged muscle tissue in mice with intact lymph nodes (**SFig. 7b-c**). We observed a dramatic and significant decrease in several lymphocyte populations including total T cells (CD4+, CD8+ and DN T cells; **Fig.3 d**), GC-like B cells (**Fig.3 e**), and importantly plasma cells (**Fig.3 f-g**) across treatments in xLN mice (**Fig.3 d-g**). Collectively, our data suggests that without lymphocyte recruitment from draining lymph nodes and retention within damaged tissue, local plasma cell differentiation is impeded.

Given the heterogeneity in lymphocyte recruitment after VML at 7dpi (**Fig.1a**) and ectopic plasma cell generation (**Fig.2c**), we hypothesized that a peripheral B cell helper T cell subset must be differentially recruited upon PE_tx_ and promote a local plasma cell response. In order to identify these cells, we performed sub-clustering of muscle infiltrating αβT cells at 7 (**Fig.3h**) and 21 dpi (**Fig.3i**).

In agreement with previous reports, ECM enriched canonical CD4+ FoxP3+ regulatory T cells, and a previously described T follicular regulatory subset^34^ (4PD-1hi T follicular regulatory; **SFig. 7d**) expressing Th2/Treg and Tfh markers (*Il4, Il21, Bcl6, Maf, Pdcd1*). This persisted to 21 days post injury showing a strong canonical regulatory and type-2 phenotype characterized by high IL-4 and IL-10 induction by ECM scaffolds (**Fig.3i**).

In contrast, T cells of both CD4 and CD8 lineage with a naïve-like phenotype (*Ccr7, Sell, Lef1,* and low *Cd44* and *Fas*) were enriched in PE at 7dpi. Further, in addition to enrichment of γδ17 T cells, a double negative αβT cell subset expressing both Th17 and follicular helper cell markers (*Rorc, Il22, Ccr6, Maf, Cd40lg, Icos,* and *Fas* while lacking *Il17a*), which have been implicated in IgA production^35,36^ was also enriched in the muscle upon PE_tx_ (**Fig.3j-k**). Notably, we did not find any T cell cluster enriched for classical B cell helper cytokines *Cxcl13* and *Il21* (**Fig.3 j-k**) that was associated with tissue plasma cell differentiation in the PE group. As such, it is possible that one or multiple peripheral follicular helper T cell subsets utilize *Cxcl13* or *Il21* independent pathways to facilitate plasma cell differentiation within injured tissue.

When evaluating potential contribution of CD4 and CD8a mediated help in maturation of plasma cells, genetic deletion of CD4 (*Cd4^-/-^*) or CD8a (*Cd8a^-/-^*) co-receptors led to a reduction of GC-like B cells and plasma cells both in the muscle and draining lymph nodes without total abrogation (**SFig. 7e-f**). Without either CD4 or CD8a co-receptor, the number of GC-like B cells in tissue reduced significantly, suggesting that cells bearing CD4 or CD8a co-receptors contribute to ectopic germinal center formation (**SFig. 7g**). Additionally, within damaged tissue, both CD4 and CD8 T cells contributed *Lta* and *Ltb* and TLS supporting chemokines^37^ (**SFig. 7h**). By tracking paired T cell receptor CDR3 sequences on individual T cells, we found several clones that were shared between lymph nodes and muscle indicating translocation, and further, a significant number of clones that had expanded within the muscle by 21 dpi (**SFig. 7i**). Lastly, a TLS associated gene signature (*Ccr7, Ccl3, Ccl4, Ltb, Ccl2, Fut7, Fut4*) was upregulated in damaged tissue with PE by 21 dpi (**SFig. 7j**). Together, our data suggests that both CD4 and CD8 T cells translocate to injured tissue and support TLS formation.

### Fas restricts plasma cell differentiation in TLS

Accumulation of naïve T cells to non-lymphoid organs is associated with autoimmunity^38,39^. Specifically, CCR7 and CD62L expressing naïve-like T cells are known to aggregate within non-lymphoid tissues and form tertiary lymphoid structures^40,41^. Among T cell subsets recruited to the muscle at 7dpi, we identified two clusters (CD4+ and CD8+) that bore markers of naïve T cells (*Sell*, *Tcf7*, *Lef1, Ccr7*) that also expressed low levels of *Fas* and *Cd44.* We identified these cells in the muscle as CD44+ Fas+ BCL6+ T cells expressing low levels of CD8a, ICOS and PD-1 (**Fig.4a-b**). The frequency of these cells was significantly higher in tissue with PE (**Fig.4c**). Additionally, the frequency and abundance of these cells in injured tissue depended on *Batf3* (**Fig.4d-e**). In addition to *Fas* expressing T cells, we also identified *Fas* expressing dendritic cells that were enriched by PE_tx_ at 7dpi (**Fig.4f-g**).

While Ly49+ CD8/NKT-like cells in tissue expressed markers associated with a cytotoxic program (*Nkg7, Klrb1c, Ctsw, Klrc1*), they did not express granzymes or perforin (*Gzmb, Gzma, Gzmk or Prf1)* and instead only expressed *Fasl* (**Fig.4h**). Additionally, both *Batf3^-/-^* and *Cd8a^-/-^* mice showed elevated CD11b-CD11c-B220+ B cells (**Fig.4i**), plasma cell counts (**Fig.4j**) and frequency of conversion (**Fig.4j**) upon ECM_tx_ as compared to WT mice. Together, these data suggested that Ly49+ CD8/NKT cells are an important contributor to Fas-Fasl mediated regulation in damaged tissue. To test whether Ly49+ CD8/NKT-like cells can target *Fas* expressing T cells, we isolated CD62L+CCR7+ T cells from the muscle of mice which received PE_tx_ at 7dpi. Concurrently, we isolated Ly49+ CD8/NKT-like cells from the muscle of mice which received ECM_tx_ at 7dpi. We expanded these cells *in vitro* for 7 days and then co-cultured for 16 hours. We stained all cells with Annexin-V and 7-AAD to detect apoptotic and dead cells. We observed enhanced elimination in T cells that were co-cultured with Ly49+ T cells indicating a capacity for T cell elimination (**Fig.4k**).

To investigate the downstream effects of defective Fas on plasma cells in tissue, we utilized B6.*Fas^lpr^*mice. Upon VML and PE_tx_, these mice showed significantly enhanced plasma cell differentiation in the muscle albeit at significantly lower numbers (**Fig. 4l-n**). Plasma cell conversion and total counts in draining lymph nodes were elevated compared to WT across treatment groups (**SFig. 7k**). Since we saw a striking difference in plasma cell conversion within tissue in *Fas^lpr^* mice, in addition to T cell translocation from draining lymph nodes in our lymphadenectomy experiment, we hypothesized that Tfh would escape regulation in mice depleted of Ly49+ CD8/NKT-like cells or mice lacking Ly49+ CD8/NKT-like activation. Indeed, CD4+ CXCR5+ PD-1+ Tfh cell counts in the draining lymph nodes of mice across treatments in *Cd8a^-/-^* and *Batf3^-/-^* were significantly elevated (**Fig.4o**). This was recapitulated by Fas perturbation alone suggesting that restricting T cell help to B cells is a key step in regulating autoimmunity triggered by tissue damage. Although plasma cell differentiation was not completely abrogated in *Cd4^-/-^*mice suggesting additional helper mechanisms, this finding that canonical Tfh were selectively expanded in *Batf3^-/-^*, *Cd8a^-/-^* and Fas^Lpr^ mice suggest that CD8/NKT-like cells normally restrain conventional T cell help. The increased plasma cell conversion in *Fas^lpr^* mice led to significantly elevated serum levels of IgA and IgG2a (**SFig. 7l**).

### Microplastics promote expansion of tissue antigen specific CDR3 sequences in type-17 T cells and B cells

In addition to the CDR3 sequences on CD8/NKT-like cells (**Fig.1e**), we evaluated clonal expansion and self-reactivity of muscle infiltrating T and B cells at 7 and 21 dpi. (**Fig. 5a-e**). Early in response to injury there was minimal but detectable expansion of CDR3s in several T cell subclusters (**Fig. 5a**). Of those that had expanded CDR3s, we found that several clusters shared the same expanded CDR3 which had a TRA sequence of CVVGDRGSALGRLHF. Through VDJdb we found that this TRA CDR3 is associated with insulin-reactive T cells. When continuing to 21 days post-injury we saw a stronger oligoclonal expansion in CD4+ T cell clusters including canonical CD4 Tregs, 4PD1hi/TFR, CD4 TCM, and Th2 T cells (**Fig. 5b and SFig. 7d**). When instead evaluating expansion of unique clones, we found that two populations had robust expansions of a single clone, the exhausted CD8 cluster and a *Klra5*+ CD8/NKT-like cluster (**Fig. 5c**), with the latter sharing transcriptional phenotypes associated with “CD8 Regulatory T cells” previously reported in controlling autoimmunity^19^. While the exhausted CD8 TRB CDR3 did not have a known antigen, the *Klra5*+ CD8/NKT-like subcluster had the same CVVGDRGSALGRLHF insulin-reactive sequence as several 7-day cell clusters. Given this difference and specifically the TRA expansion, we evaluated overall TRB versus TRA expansion and found that there was also a significant expansion of TRA in an innate-like Th17 population enriched by polyethylene (**Fig. 5d**). As with the prior analyses, this same CVVGDRGSALGRLHF sequence reappeared. When analyzing overall distribution of CVVGDRGSALGRLHF bearing cells within ECM_tx_ and PE_tx_ mice, we find a strong preference for the immunoregulatory-like population (blue) in ECM while PE showed a strong enrichment in the innate-like Th17 cluster (**Fig. 5e**).

In addition to the local muscle microenvironment, we sequenced TCR CDR3s in the draining inguinal lymph node (**Fig. 5f-g**). Evaluating TRA CDR3s that were expanded more than 10 copies in the inguinal lymph node, most sequences shared a similar CA-starting sequence with one expanded CDR3 sequence standing alone in sequence similarity to other expanded CDR3s, which was the same CVVGDRGSALGRLHF insulin reactive clone (**Fig. 5f**). As with the muscle tissue, there was a preference for this CDR3 to be enriched in type-17 polarized cells for PE_tx_ mice in comparison to ECM_tx_ mice (**Fig. 5g**). Indeed, when evaluating reactivity against insulin, we found induction of circulating autoimmune antibodies (anti-insulin IgG) at 21 days post-injury that persisted out to 4 months post-injury (**Fig. 5h**). Additionally, insulin specific circulating IgA was significantly elevated in PE at 21 days post injury (**Fig.5i**).

When evaluating CDR3 sequences of these early infiltrating B cells in WT C57BL6 mice, we found both shared phenotypes and divergent enrichment of amino acids in their B cell receptors (**SFig. 8a**) across treatment groups. In all treatment groups the most abundant CDR3 was CALWYSNHWVF that is associated with Smith Antigen (Sm)^42^ reactivity, an snRNP and lupus marker (**Fig. 5j-k**). The total number of anti-Sm BCR was highest in the muscle of PE_tx_ mice (**Fig. 5j**) and was restricted to lymph nodes of ECM_tx_ mice (**Fig. 5k**). In addition, we also found several sequences that were shared between a control injury and PE_tx_ (**Fig. 5l-m**) and uniquely PE-specific clones (**Fig. 5n**) including some with the sequence CQQYYSYPPTF in the CDR3 (**Fig.5n**) and that has been reported to result in binding to LPS, insulin, and dsDNA in autoimmune conditions^43^ while not binding microbiome antigens^44^. Circulating IgG antibodies in NOD mice showed robust banding against uninjured muscle lysates (**Fig. 5o,q**) whereas in C57BL/6 mice, PE treatment significantly increased circulating muscle antigen specific IgG (**Fig.5p,r**) supporting a polyclonal autoimmune response.

We previously noted that PE induced elevation and persistence of insulin-reactive IgA (**Fig. 5i**). To evaluate potential clinical relevance, we mined a scRNAseq dataset of human type 2 diabetes (insulin insensitivity) to evaluate alterations in immune cells in the peripheral blood. Here, we found a decrease in CD8+NKT/MAIT-like cells that expressed KIRs, XCL1/2, and HELIOS (**Fig.5s**). These cells expressed lower levels of transcripts for FASLG along with XCL2 in T2D patients suggesting impaired killing function (**Fig.5t**). Interestingly, the most enriched TCR was a CMV-reactive CVVSDRGSTLGRLYF, which is 3 amino acids away from the insulin reactive clone observed in our LCMV-naïve mice (**Fig.5u**). Lastly, to investigate the relevance of KIR+ CD8/NK T cells to TLS in humans we reanalyzed a spatial transcriptomic dataset from humans with diabetic kidney disease^45^. Among spatial sections where KIR+ cells were detected, we analyzed prevalence of dendritic cell, helper T cell and B cell markers (**SFig. 8b-c**) and performed nearest neighbor analysis. We observed that KIR+ cells surrounded TLS-like niches. Interestingly, the mean Euclidean distance from *Itgax, Fas* and *Rorc* was significantly lower as compared to *Cxcl13* or *Il21* (**SFig. 8d**). Taken together, this suggests that KIR+ T cells are recruited to inflamed and damaged tissue in humans and might have distinct targets in lymph nodes and TLS.

## DISCUSSION

Traumatic injury results in release of self-antigen and persistent inflammation that rarely lead to persistent autoimmunity. Following tissue damage, we observed that lymphocytes organize into TLS and give rise to plasma cell locally. Further, we describe an immune editing regulatory pathway that limits the plasma cell response within TLS, averting autoimmunity in times of acute, significant inflammation and tissue damage. Rapid accumulation of cDC1s that we previously showed was mediated by canonical NK cells^10^ supports the activation of a Fas-dependent Ly49+ CD8/NKT-like population that prunes autoreactive immune cells. Self-antigen can be hydrophobically trapped by high surface area contaminants such as microplastics^46^. Coupled with persistent inflammation of injured tissue with fibrotic material and the ensuing foreign body response, this creates an environment poised for self-reactivity. Indeed, we found that these microplastics give rise to local germinal center reactions after tissue damage. This promotes ectopic differentiation of autoreactive plasma cells driven by draining lymph node derived T cells. In agreement with previous reports, we saw that TLS can drive both antigen specific^47^ as well as polyclonal^48^ T cell responses. These findings extend current models of immune tolerance after tissue damage that have largely focused on FoxP3+ regulatory T cells downstream of acute injury. Whereas previous studies have implicated Ly49+/KIR+ CD8 T cells in maintenance of peripheral tolerance during chronic autoimmunity^19,22,49^, our findings identify a functionally similar population that operates during acute tissue injury and connects cDC1-dependent regulation with tertiary lymphoid biology.

One unexpected observation was the expansion of an insulin-specific T cell receptor clone in this CD8/NKT-like population when injuries were treated with a material used to promote wound healing. Conversely, this insulin-specific clone was differentially expanded in a pathogenic Th17-like population in microplastic contaminated wounds and was associated with the development of insulin reactive IgG and IgA. With the known incidence of insulin insensitivity in trauma patients with chronic critical illness (CCI) and a recent report showing IL-17 signatures in the periphery as predictive of CCI^50^, these data suggest the local wound microenvironment as a critical mediator of post-injury immune tolerance.

This study has several limitations that are important to consider while interpreting the data. Firstly, there are no specific knockout models of Ly49+ CD8/NKT-like or CD8 regulatory T cells. These cells express Ly49 proteins, whose genes are known to be variegated, stochastically expressed and share functions, and therefore difficult to delete^51,52^. Their developmental relationship to NKT Cells, canonical CD8 T cells, and other innate-like lymphocytes is unknown. Although CDR3 sequencing of the T and B cell repertoire revealed a polyclonal response, the complete antigenic landscape of the TLS response is unknown. While our studies show that shared self-antigen specificity between T and B cells led to insulin specific autoantibodies after VML, we have not uncovered the mechanism of B cell help within damaged tissue, owing to low sequencing depth or abundance of such cells. Finally, while polyethylene was selected due to its ubiquitous use in medical devices and environmental materials, conservation of the TLS phenomenon dependent upon plastic chemistries is yet to be determined.

We propose a model in which tissue injury generates autoreactive lymphocytes that are normally constrained by a cDC1-dependent pathway inducing Ly49+ CD8/NKT-like immune editing via Fas-FasL interactions. While injury-associated TLS would be beneficial in the context of persistent inflammation due to infection, sterile microplastics reduce engagement of the cDC1-CD8/NKT pathway leading to TLS formation and plasma cell accumulation in damaged tissue. Together, these data provide a mechanistic framework for peripheral tolerance regulation in acute traumatic injury settings that connects tissue damage, wound contamination, and a potential relationship to downstream autoimmunity.

## MATERIALS & METHODS

### Preparation of biomaterials for VML

Biomaterials used as implants in mice were prepared as reported previously^10,53,54^. Small intestines obtained from 5 to 6-month-old American Yorkshire pigs were washed with water to remove feces and mechanically scraped to remove cell and mucus layers. The sub-mucosa (SIS) was then cut into 1 to 2-inch segments, washed in sterile deionized water and incubated in 10% Antibiotic-Antimycotic solution overnight at room temperature while stirring to remove any remaining fecal matter and mucosal tissue. Following incubation, SIS was washed with sterile deionized water and incubated at room temperature for 30 minutes in a solution containing 0.1% peracetic acid and 4% ethanol in sterile deionized water. Following this, SIS was subjected to alternating washes in filter sterilized tap water and 1x PBS until pH neutralization. Lastly, SIS was lyophilized for 48 hours and cryogenically milled to obtain a fine powder which was rehydrated to 200 mg/mL with a paste-like consistency before injecting at the site of implantation through a sterile syringe. Polyethylene (PE) particles of 150 µm diameter were subjected to shortwave UV-irradiation for 40 minutes, resuspended in 70% ethanol overnight, transferred to Eppendorf tubes and air dried in a sterile biosafety cabinet.

### Volumetric muscle loss and lymphadenectomy procedures

All mice received bilateral volumetric muscle loss procedures unless noted otherwise. One to three days prior to surgery 6–8-week-old mice C57BL/6 WT (Jackson Laboratory stock # 000664), *Cd8a^-/-^* (JAX Stock # 00266532), *Batf3^-/-^* (JAX Stock # 01375533), B6. *Fas^lpr^* (JAX Stock # 000482), B6.*Aire^-/-^* (JAX Stock # 00474334), *Cd4^-/-^* (JAX Stock #002663) NOD and NOD.*Aire^-/-^* (Taconic Biosciences) mice were cleared of hind limb and abdominal fur by shaving with electric razor followed by depilatory cream application. Anesthesia was first induced in a chamber supplied with 4% isoflurane and maintained with a nosecone supplied with 2% isoflurane supplemented with oxygen at a flow rate of 200 mL/min. Skin overlying the quadriceps was sterilized three times with betadine and isopropanol alternatingly in a circular pattern working from the incision site outward. Following this, a 1cm surgical incision into the skin and fascia was made to isolate the quadriceps muscle and a 3mm defect was created centrally using sterile surgical scissors. The wound was then backfilled with either ECM, PE or no biomaterial as a control. NOD WT and *Aire^-/-^* mice were used following identical procedures. All mice were monitored until reported endpoints. The protocols (NIH Clinical Center ACUC protocol NIBIB23-01 and NIAD protocol LCIM 14E) were reviewed approved by the National Institutes of Health Scientific Review and Animal Care and Use Committees. The animal facilities are and tested annually for ECTRO, EDIM, LCMV, MAV1, MAV2, MHV, MPV, MVM, MYCO, NS-1, POLY, PVM, REO3, SENDAI and TMEV pathogens. In the facility for most studies (all except NOD and NOD. *Aire^-/-^*) no positive sentinel reports have occurred in the last 11 + years except for non-pathogenic flagellates. Helicobacter is not tested for, MNV is excluded in the facility. For NOD and NOD. *Aire^-/-^* mice, MNV and heliobacter are suspected in the facility.

### Isolating dendritic cells and lymphocytes

C57BL/6 WT mice which received biomaterials and control injuries were euthanized at 7 days post-injury via carbon dioxide asphyxiation followed by cervical dislocation. Quadriceps along with biomaterials were dissected, minced, digested in a buffer comprising 0.25 mg/mL Liberase TM and 0.2 mg/mL DNAse I in serum-free Dulbecco’s Modified Eagle Medium (DMEM) for 1 hour, and passed through a 70 µm cell strainer. Inguinal lymph nodes were mashed and passed through a 70 µm cell strainer and washed with 1x PBS. The resulting single cell suspensions were stained with 7-AAD for 10 minutes and a surface marker cocktail to isolate dendritic cells from muscle tissue and lymphocytes from muscle tissue and inguinal lymph nodes respectively. Antibodies identifying surface markers for DC, B and T cell isolation are in Supplemental File S2. Cells were sorted using a BD FACSMelody sorter into a buffer comprising 50% FBS and 25mM HEPES in 1x PBS. For single cell RNA sequencing analysis, cells were centrifuged at 400 xg at 4 C for 5 minutes, resuspended in Recovery™ Cell Culture Freezing Medium (ThermoFisher Scientific), frozen overnight at –80°C in a slow cooling apparatus and transferred to a liquid nitrogen tank until further analysis. Cells were shipped on dry ice to Azenta Life Sciences for single cell RNAseq.

### Flow cytometry and data analysis

Whole blood leukocytes, quadriceps muscle, and inguinal lymph nodes (ILN) were obtained from mice at predetermined endpoints using standard dissection and isolation procedures. Muscle tissue was minced and either digested in 0.05% Collagenase IV and 0.2 mg/mL DNAse I to stain for dendritic cell markers or directly mechanically dissociated on a 70 µm cell strainer and washed with a buffer comprising 1% BSA and 2 mM EDTA to stain for enzyme sensitive lymphocyte markers. Peripheral blood was collected through submandibular bleeding in 5mM EDTA in 1x PBS and red blood cells (RBC) were lysed using eBioscience 1x RBC Lysis buffer (Multi-species) before analysis. Lymph nodes were mechanically dissociated through a 70 µm cell strainer and washed with a buffer comprising 1% BSA and 2 mM EDTA (wash buffer). Resulting single cell suspensions were washed in 1x PBS and resuspended in 200 µL of Live/Dead Blue (1:1000 dilution) for 20 minutes at 4°C. Cells were then washed, resuspended in 50 µL of solution containing TruStain Monocyte block (1:10 dilution) and TruStain Fc block (1:100 dilution). Separate panels for dendritic cell markers and lymphocyte markers were used to analyze samples. Using these panels, cells were stained for 1 hour at 4°C, washed in wash buffer at 400 xg for 5 minutes at 4°C and fixed using True-Nuclear Transcription Factor kit (Biolegend). Lymphocytes were washed in a permeabilization buffer (True-Nuclear Transcription Factor kit (Biolegend)) at 400 xg for 5 minutes at room temperature and stained overnight for intracellular markers at 4°C. Cells were washed at 400 xg for 5 minutes at 4°C and resuspended in wash buffer prior to analysis on a 5 laser Cytek Aurora spectral flow cytometer. Data were unmixed via single-spectra controls using SpectroFlo (v3.2.1) then unmixed data were analyzed via FlowJo (v10.10.0). Resulting data were analyzed on GraphPad Prism v10.3.1.

### Single cell RNA sequencing raw data processing, analysis and visualization

Sequenced reads were aligned, and feature-barcode matrices were generated using Cell Ranger. Matrices were read into R and the DropletsUtils package was used to filter out empty droplets and generate final filtered counts for analysis. The Seurat package (v4.3.0) was used to filter out cells containing > 20% mitochondrial or ribosomal counts for each treatment group. Finally, samples from all treatment groups were merged to create a unified Seurat object. Following the standard Seurat pipeline, Uniform Manifold Approximation and Projection (UMAP) clusters were generated after logarithmic normalization (LogCP10K), data scaling and top 2000 features selected using variance stabilizing transform method for principal component analysis (PCA) retaining the top 50 principal components for computing nearest neighbors. Cluster identity was assigned using either canonical or published literature. Latent time analysis was run in Python v3.8.18 using the “SpatialCorr” and “Scanpy” libraries. For mined mouse data, GSE20028922 and GSE14247121 were retrieved from GEO and subclustered prior to display of CD8/NKT-like gene signature on T cell sub-cluster. For mined human scRNAseq data, GSE14370423 of human skeletal muscle tissue was subset to evaluate CD3+ T cells. Within this subset, cells expressing genes encoding KIRs and IKZF2 were displayed using GraphPad Prism v10.3.1.

### T and B cell receptor sequencing data analysis and visualization

T and B cell receptor data was analyzed using the “cellranger vdj” pipeline. FASTQ files were input via command line to run “cellranger vdj” and filtered and annotated outputs were used for downstream analysis. For visualizing gene usage, the Vβ segments were filtered and displayed using a pie chart. Clonotypes generated using cellranger were analyzed using the “immunarch” package in R (4.2.1). Clonotypes were tracked between muscle and lymph node datasets using the “trackClonotypes” function based on amino acid sequence matching. CDR3 length and hydrophobicity scoring were analyzed using Excel with relevant CDR3 sequence lengths as stated in the text, then data were displayed using GraphPad Prism v10.3.1. Logo plots were generating using WebLogo (UC Berkeley). CDR3 length and hydrophobicity scoring were analyzed using Excel with relevant CDR3 sequence lengths as stated in the text, then data were displayed using GraphPad Prism v10.3.1. Potential antigens were identified using https://VDJdb.org searching for appropriate TRA or TRB CDR3 sequences.

### Spatial transcriptomics data analysis and visualization

Five (5) μm cross sections of quadriceps were taken from mice which received VML and biomaterial implants at 1 week post injury. Sections were H&E stained and imaged using a Leica brightfield microscope for mapping transcripts to tissue location. Spatial transcriptomic sequencing was carried out using a 10X Visium instrument at the NICHD molecular genomics core facility. The standard Space Ranger workflow was used to generate gene expression matrices using the mm10 reference transcriptome. Outputs were subsequently imported into an R Studio workspace to run the Seurat pipeline for analyzing spatial datasets. Counts were normalized using the function “SCTransform” and gene expression was visualized using the “SpatialFeaturePlot” function. To plot signatures from single cell RNA sequencing datasets, references were normalized using the “SCTransform” function and an anchor-based integration method was used to generate prediction scores for each spatial spot. Additionally, we also generated module scores for gene sets containing enriched genes in specific clusters and visualized the score using “SpatialFeaturePlot”.

### Bulk RNA sequencing data analysis and visualization

Bulk RNA sequencing data analysis was performed as previously reported^54,55^. Briefly, whole quadriceps muscles were flash-frozen on dry ice for bulk RNA isolation and sequencing to Azenta GeneWiz. Data analysis was performed on the paired end reads from 12 independent samples. Pseudoaligment was performed using Kallisto to map reads to a publicly available reference transcriptome (Ensemble GRCm39). The Ensemble Mus musculus v79 annotation database mapped transcripts to gene IDs. EdgeR package was used to determine counts per million and subsequently filtered by counts per million > 1 for at least 3 samples and log2 transformed. Data was normalized using the Trimmed Mean of M-values method. Differential expression was visualized using standard functions to generate fold change plots for each pairwise comparison. Gene set enrichment analysis (GSEA) was carried out in R using the “GSEABase” package and molecular signatures gene set collection.

### Protein isolation and bone marrow-derived cDC1 generation

Decellularized ECM and muscle tissue were homogenized at 5000 rpm for 30 seconds using a handheld homogenizer (Fisherbrand homogenizer 850) in RIPA buffer (ThermoFisher) containing protease inhibitors (Pierce, ThermoFisher). Isolates were diluted, resuspended in sterile deionized water and quantified using Pierce BCA assay following manufacturer’s instructions. Protein solutions were stored at –20 C until further use. Bone marrow cells were isolated from the femur bones of healthy 8-week-old C57BL/6 mice. Cells were cultured for 9 days in media containing GM-CSF (5 ng/mL), FLT3L (150 ng/mL) in complete RPMI 1640 medium (10% FBS, 1% Penicillin/Streptomycin) with glutamine as described elsewhere^56^. Weakly adhered and floating cells were collected, and media was replenished on days 3 and 6 of culture. Resulting cells were matured overnight by lipopolysaccharide (LPS) stimulation (50 ng/mL) and loaded with protein isolated from ECM and muscle tissue (1 µg/mL).

### In vitro T cell stimulation and proliferation

Ly49+ CD8+ T cells were isolated flow cytometrically as described above. T cells were washed at 400 xg for 5 minutes at 4 C and resuspended in 1x PBS containing cell trace violet stain (1:1000 dilution) and incubated in a water bath at 37 C for 20 minutes. T cells were washed in complete media and resuspended in complete RPMI 1640 medium (10% FBS, 1% Penicillin/Streptomycin) supplemented with IL-2 (1ng/mL), GM-CSF (5 ng/mL), FLT3L (100 ng/mL) and 50 µM β-mercaptoethanol. T cells were cultured with induced cDC1s loaded with either protein isolated from ECM. A 1:2 ratio of APCs:T cells was used and cell culture was carried out for 3 days prior to analysis on Cytek Aurora flow cytometer.

### In vitro apoptosis and cell death detection

CD4-Ly49+ T cells and CCR7+CD62L+ T cells were isolated at 7 days post-injury from muscle tissue of mice which received ECM_tx_ and PE_tx_ respectively. Cells were stimulated with soluble CD3ε antigen (1 µg/mL) for 24 hours in the presence of recombinant IL-2. Following stimulation, cells were washed with 1x PBS and replated in complete RPMI 1640 media containing recombinant IL-2 (10 ng/mL) for 6 additional days. Media was replenished every 48 hours. CCR7+CD62L+ T cells were then co-cultured with CD4-Ly49+ T cells for 16 hours. Cells were pelleted, washed and stained with conjugated antibodies against Annexin-V (eFluor-450), CD62L(APC-Cy7), CCR7(APC), Ly49(PE) in Annexin-V binding buffer for 30 minutes at room temperature followed by washing and staining with 7-AAD for 30 minutes at room temperature and analyzed on a Cytek aurora flow cytometer.

### Immunohistochemistry and Immunofluorescence

Human and mouse tissue samples taken were fixed in neutral buffered formalin for 48 hours prior to sequential dehydration in 70%, 80%, 95% and 100% ethanol. Samples were then cleared in xylene and paraffin embedded before sectioning at a thickness of 5 µm. Sections were heated to 60 ⁰C overnight, cooled to room temperature, rehydrated, incubated in boiling citrate antigen retrieval buffer for 20 minutes and cooled to room temperature. All sections were blocked using a solution containing 4% normal goat serum, 1% bovine serum albumin and 0.05% Tween-20 in 1x PBS for 1 hour. Following this, samples were incubated with primary antibodies (Rat anti Mouse B220; BioLegend and Rabbit anti Mouse CD3e; Abcam) overnight at 4 ⁰C in a humidity chamber. After three washes in 0.05% Tween-20 in 1x PBS, samples were incubated with secondary antibodies (Donkey anti Rat Alexa Fluor-594; Invitrogen and Goat anti Rabbit Alexa Fluor-488; Invitrogen) for 90 minutes at room temperature. Autofluorescence was quenched in all samples using Vector True View autofluorescence quenching kit (Vector Labs) following manufacturer’s instructions. Finally, samples were counter stained with DAPI, washed in DI water, mounted and imaged using an EVOS M5000 microscope. Images were adjusted for brightness and contrast in FIJI (ImageJ2, Version 2.14.0/1.54f) using primary delete controls to set thresholds.

### ELISA analysis

Peripheral blood from mice undergoing VML was collected by submandibular bleeds on the day prior to surgery and at indicated timepoints after surgery. Subsequently, serum was isolated by centrifugation and stored at -20 C for short term and -80 C for long term storage until analysis. Sera were diluted 1:10,000; 1:500,000; 1:20,000 or 1:100 to measure total circulating IgA, IgM, IgG3 and IgE respectively using uncoated ELISA kits (Thermo Fisher) using manufacturer instructions. Sigmoidal four parameter logistic curve fitting was used to interpolate a standard curve and OD values obtained from samples using GraphPad Prism v10.4.1.

### Statistical analysis and quantification

Statistics were computed in R (v4.2.1) and GraphPad Prism (v10.4.0). Bulk RNA-sequencing data was analyzed using empirical Bayes statistics for differential expression method using the R package “limma”. To compare various lymphocyte subsets using flow cytometry, unpaired T-tests were used to compare knock out mice to WT controls. In the co-culture assay, to compare means of the two indicated populations, a one-tailed paired T-test was used. One-way ANOVA tests with Tukey’s correction for multiple comparisons were used to compare various populations across treatment groups within WT mice and to compare populations across various knock out mice with ECM_tx_. Where indicated, normality testing was performed using the Shapiro-Wilk test. Datasets which failed normality testing were analyzed non-parametrically. Muscle samples with very low T cells yield (less than 500 cells) were omitted from analysis. None of the samples from the lymph nodes were omitted. Transcriptomic signature scores were calculated using the “AddModuleScore” function within Seurat using the genes represented in Table S1. To compare accumulation of naïve T cells in the muscle across treatment groups via flow cytometry, a mixed effects model approach was used. Gene ontology was obtained from the STRING database and gene set enrichment analysis was performed using the “GSEABase” and “msigdbr” packages within R. For correlation testing, simple linear regression analysis was done in GraphPad Prism (v10.4.0). Unless otherwise indicated, ns = p>0.05, * = p<0.05, ** = p≤0.01, *** = p≤0.001 and **** = p≤0.0001. “r” values represent Pearson’s correlation coefficient.

## Data and Code Availability

Raw sequencing data will be deposited in GEO upon acceptance of manuscript for publication. Flow cytometry raw data files will be uploaded to https://FlowRepository.org and all other data will be provided as a Source Data spreadsheet as supplemental materials. Code used for the analysis of scRNAseq data will be uploaded to GitHub. Additional figures and data are available in the **Supplementary Materials** document.

## ACKNOWLEGEMENTS

The authors are grateful to Dr. Fabio Rueda Faucz, Dr. James R Iben and Cameron Padilla at the National Institutes of Child Health and Human Development (NICHD) Molecular Genomics Core for advice on Spatial Transcriptomics, and Drs. Ronald Germain, Ethan Shevach, Angela Thornton and Matthew Wolf for helpful discussions. This work was supported by the National Institutes of Health (NIH) Intramural Research Program of the National Institute of Biomedical Imaging and Bioengineering and the National Institute of Allergy and Infectious Diseases. <u>Disclaimer</u>: The contents of this manuscript reflect the opinions of the authors solely and not the National Institutes of Health. The organizations mentioned and products used herein do not imply endorsement by the US Government.

