## Supplementary Materials for "Microplastics disrupt a cDC1-CD8/NKT regulatory circuit that restrains plasma cell accumulation after injury"

**SUPPLEMENTAL INFORMATION**

**Josyula et al.**

**Supplemental Figures 1-8**

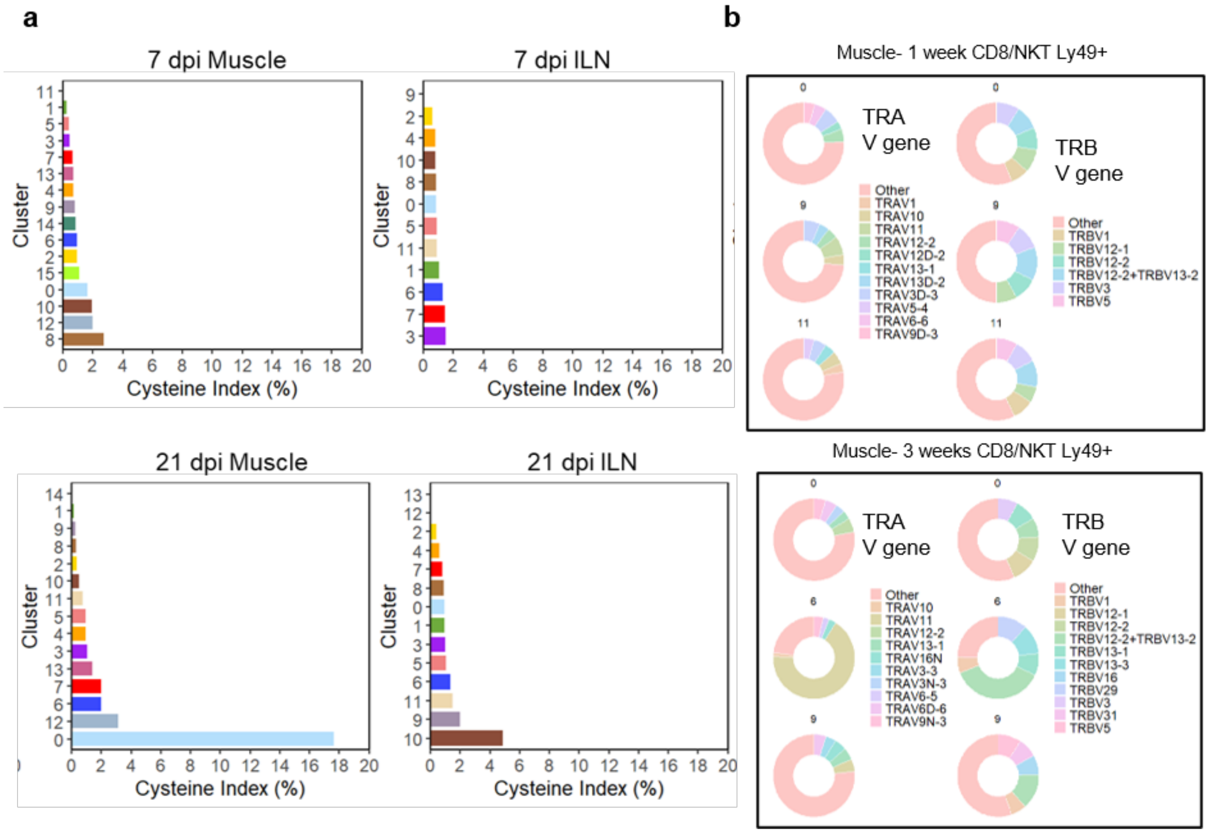


**Supplemental Fig.1 | Self reactivity and T cell receptor gene usage in CD8/NKT-like cells**. (**a**) Cysteine index as a fraction of T cells from seurat clusters shown in main figure 3. (**b**) TRA and TRB variable region gene usage in WT CD8/NKT cells expressing Ly49 receptors at indicated timepoints.

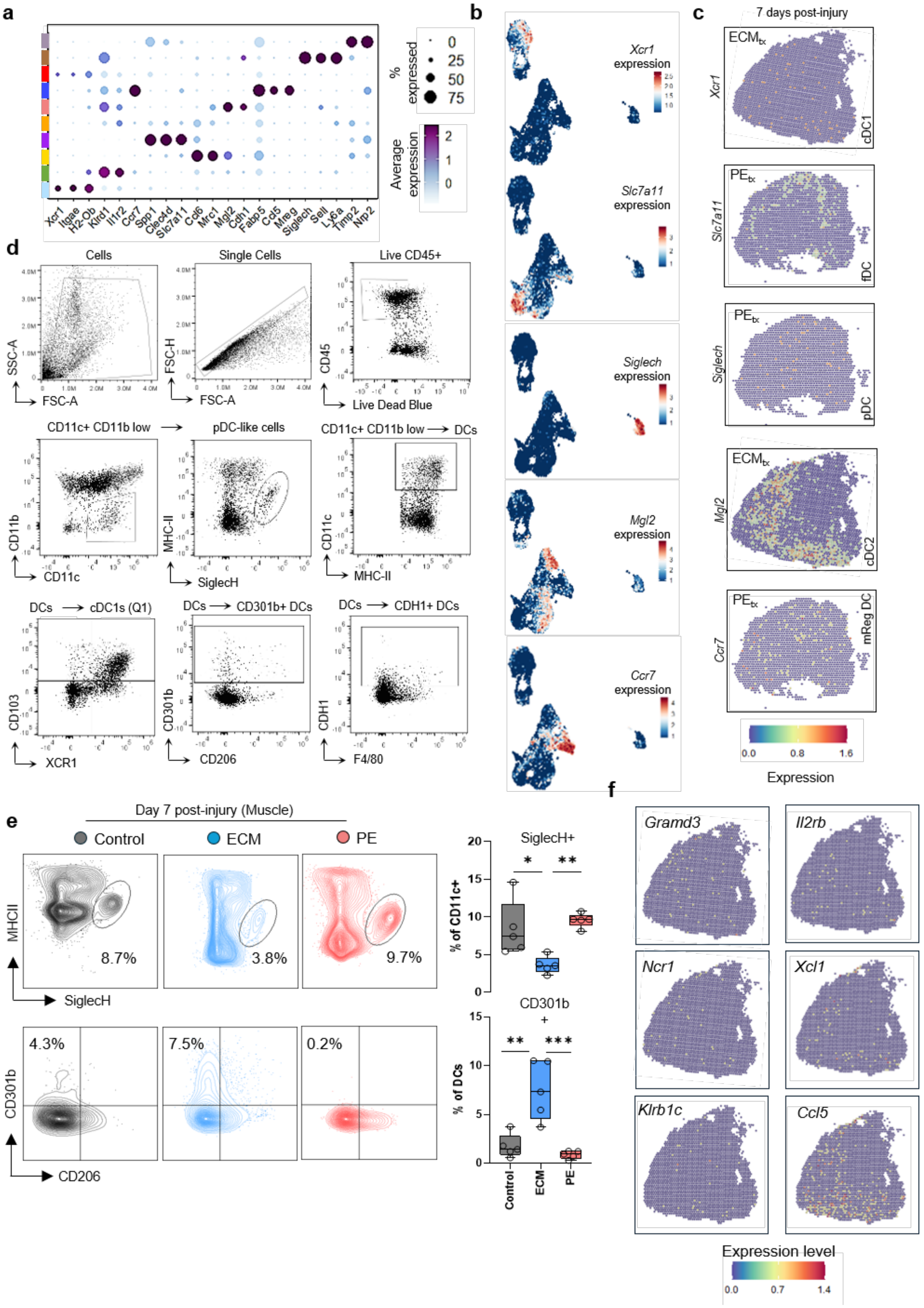

a
Xcr1
Itgae
H2-Ob
Klrd1
Il1r2
Ccr7
Spp1
Clec4d
Slc7a11
Cd6
Mrc1
Mgl2
Cdh1
Fabp5
Cd5
Mreg
Siglech
Sell
Ly6a
Timp2
Nrp2
% expressed
0
25
50
75
Average expression
2
1
0
b
Xcr1 expression
Slc7a11 expression
Siglech expression
Mgl2 expression
Ccr7 expression
c
7 days post-injury
ECMtx
Xcr1
cDC1
PEtx
Slc7a11
fDC
PEtx
Siglech
pDC
ECMtx
Mgl2
cDC2
PEtx
Ccr7
mReg DC
0.0
0.8
1.6
Expression
d
Cells
SSC-A
FSC-A
Single Cells
FSC-H
FSC-A
Live CD45+
CD45
Live Dead Blue
CD11c+ CD11b low
CD11b
CD11c
pDC-like cells
MHC-II
SiglecH
CD11c+ CD11b low → DCs
CD11c
MHC-II
DCs → cDC1s (Q1)
CD103
XCR1
DCs → CD301b+ DCs
CD301b
CD206
DCs → CDH1+ DCs
CDH1
F4/80
e
Day 7 post-injury (Muscle)
Control
ECM
PE
MHCII
SiglecH
8.7%
3.8%
9.7%
CD301b
CD206
4.3%
7.5%
0.2%
SiglecH+
% of CD11c+
CD301b+
% of DCs
Control
ECM
PE
f
Gramd3
Il2rb
Ncr1
Xcl1
Klrb1c
Ccl5
Expression level
0.0
0.7
1.4


14

**Supplemental Fig.2 | Dendritic cell recruitment to damaged tissue at 7 dpi in WT C57BL6 mice**. (**a**) Differential expression of characteristic dendritic cell markers in FACS sorted CD45+ CD11c+ MHC-II+ cells in muscle tissue at 7 dpi in dot plot and (**b**) UMAP clusters. (**c**) Spatial distribution of defining markers of dendritic cell sub-types in tissue with indicated treatments at 7 dpi. (**d**) Flow cytometry gating to identify dendritic cell fractions in injured tissue. (**e**) fractions of CD11c+ SiglecH+ plasmacytoid DCs and CD301b+ CD206- DCs across indicated treatments in damaged muscle tissue. (**f**) Spatial distribution of indicated CD8/NKT cell signature markers in mice (ECM group) at 7dpi.

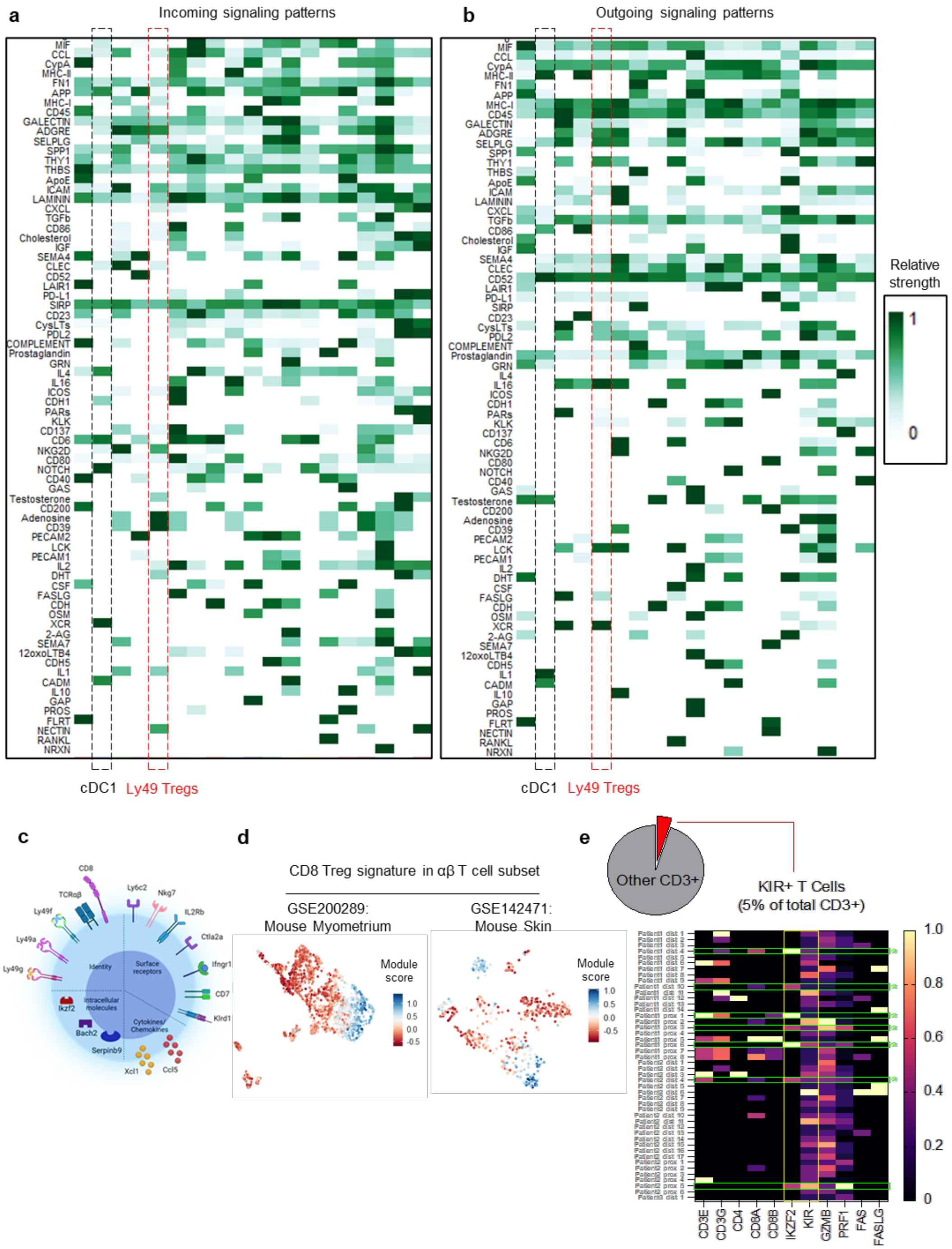


**Supplemental Fig.3 cDC1s support activation of CD8/NKT-like T cells after tissue damage in mice and humans.**(**a**) CellChat analysis showing possible communication incoming and (**b**) outgoing pathways between cDC1s and CD8/NKT-like T cells after tissue damage- analysis was performed using differentially expressed markers obtained from single cell RNA sequencing datasets introduced within this manuscript. (**c)** The transcriptomic signature of CD8/NKT-like T cells comprising *Ccl5, Ikzf2,* Ly49 (inhibitory) receptors, *Nkg7, Ly6c2, Klrd1* is present in (**d**) mouse models where myometrium (GSE20289) or skin (GSE142471) were subjected to injury. (**e**) Human analogues of characteristic markers of CD8/NKT regulatory cells were mapped onto a human single cell dataset (GSE143704) showing that CD8/NKT-like cells comprise approximately 5% of T cells in healthy donor human skeletal muscle tissue.

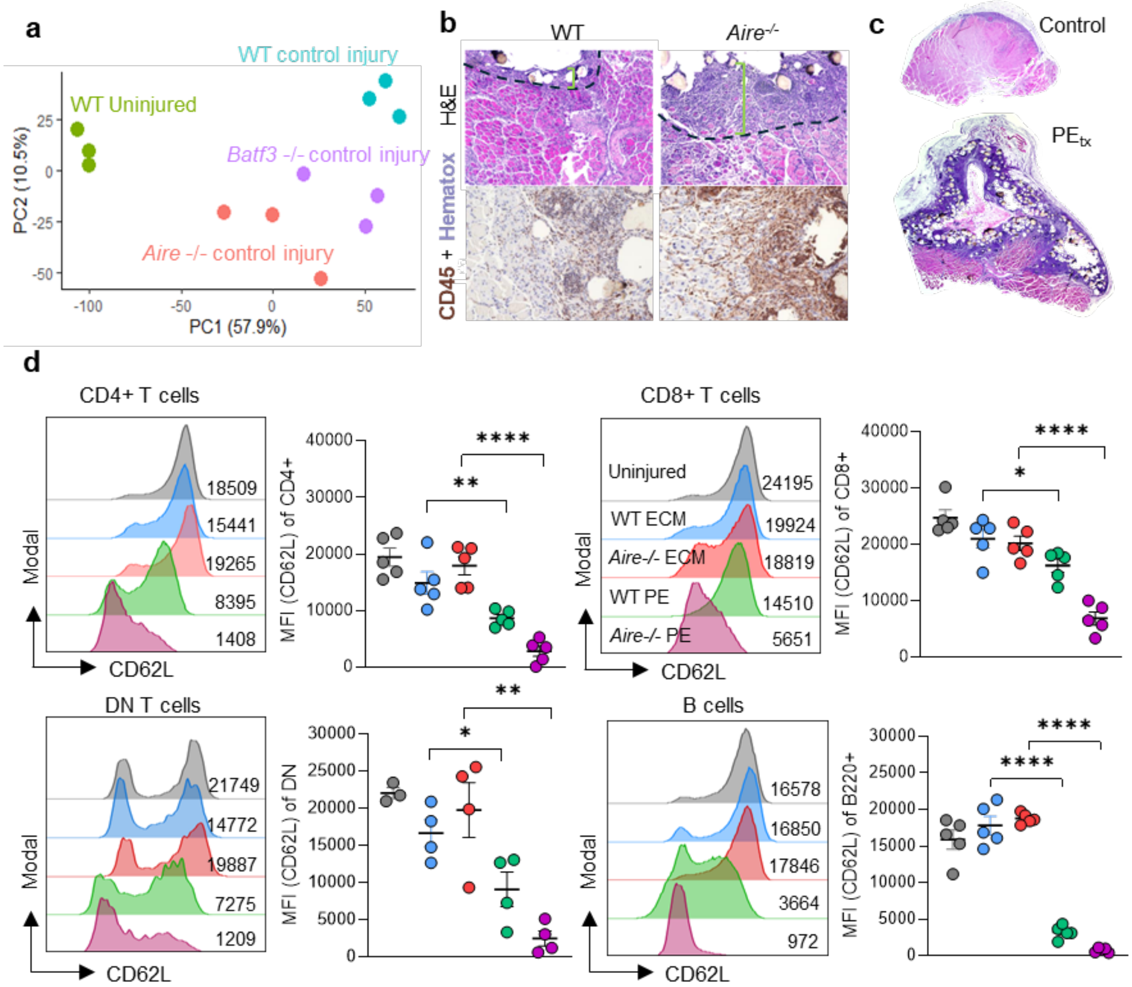


**Supplemental Fig.4 | Mice that lack cDC1s mirror and autoimmune phenotype with heightened inflammation in muscle injury when exposed to microplastics**. (**a**) PCA plot of uninjured mice, WT C57BL6, *Aire*$^{-/-}$ and *Batf3*$^{-/-}$ mice from bulk RNA sequencing performed on injured tissue at 7dpi. (**b**) Hematoxylin and eosin (H&E) and CD45 staining of WT and *Aire*$^{-/-}$ mice at 7 days post-injury (dpi) with $PE_{tx}$. (**c**) Whole cross-section of control and $PE_{tx}$ injury in *Aire*$^{-/-}$ mice. (**d**) CD62L expression in WT and *Aire*$^{-/-}$ mice on CD4+ T cells, CD8+ T cells, Double negative (DN) T cells, and B Cells in the inguinal lymph node of injured mice (7dpi) in indicated conditions. Data are mean ± standard error (SEM). Data in (**d**) are mean ± SEM in n≥3 mice.

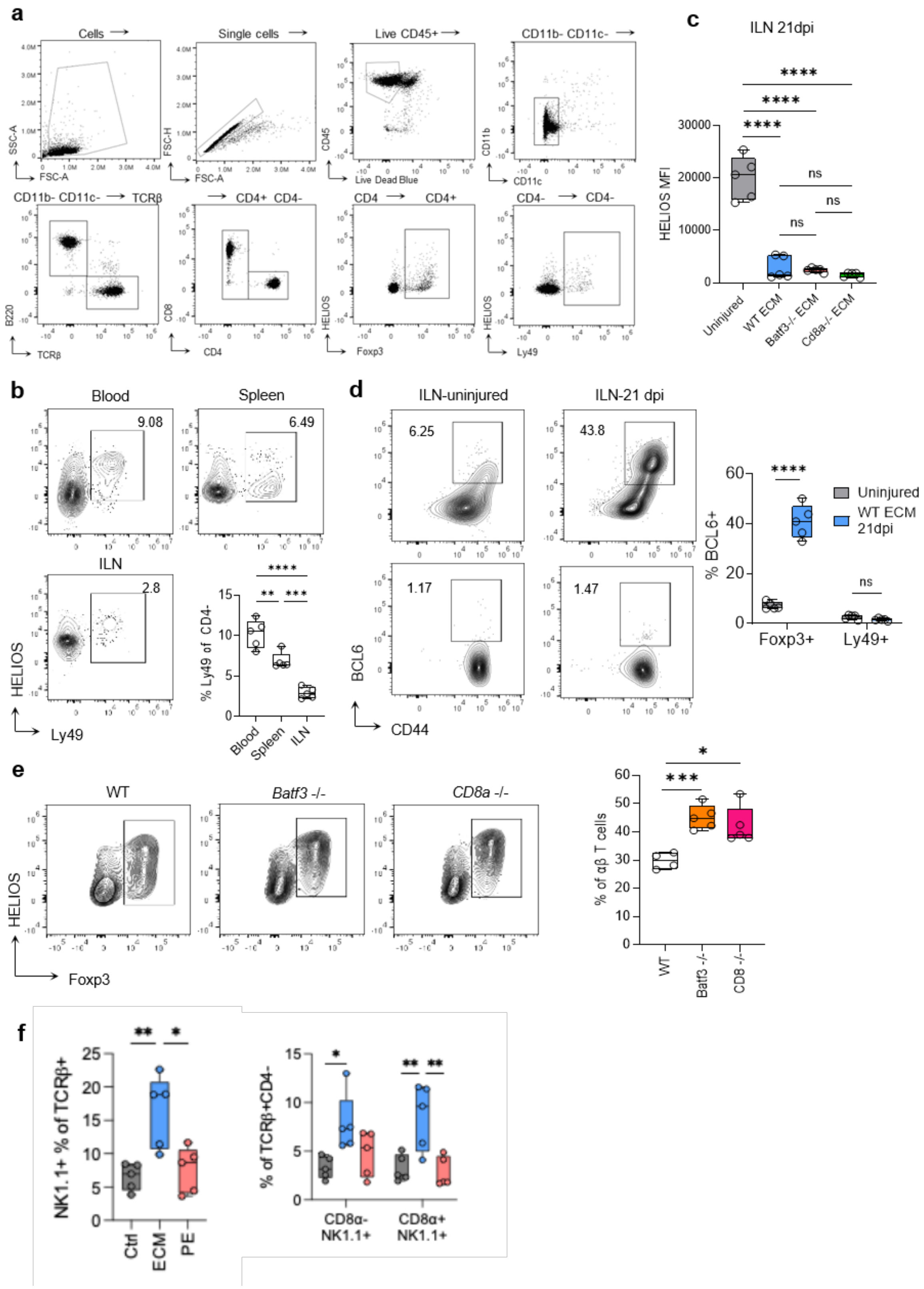


**Supplemental Fig.5 | Flow cytometry profiling of canonical CD4+ Foxp3+ regulatory T cells, CD8/NKT regulatory T cells and plasma cells.** (**a**) Flow cytometry gating to identify Foxp3+ and Ly49+ T cell populations. (**b**) Frequency of Ly49+ expressing CD4- T cells in peripheral blood, spleen and inguinal lymph nodes of uninjured WT C57BL6 mice. (**c)** Median fluorescence intensity of HELIOS in Ly49+ T cells at 21 dpi within inguinal lymph nodes of C57BL6 mice which received indicated treatments. (**d)** BCL6 expression in Ly49+ (bottom) and Foxp3+ (top) T cell subsets at 21 dpi in inguinal lymph nodes of mice which received indicated treatments. (**e**) CD4+ Foxp3+ Treg recruitment to injured muscle at 7 dpi in WT, *Batf3-/-* and *Cd8a-/-* mice. (**f**) NKT cell recruitment to injured skeletal muscle at 7dpi.

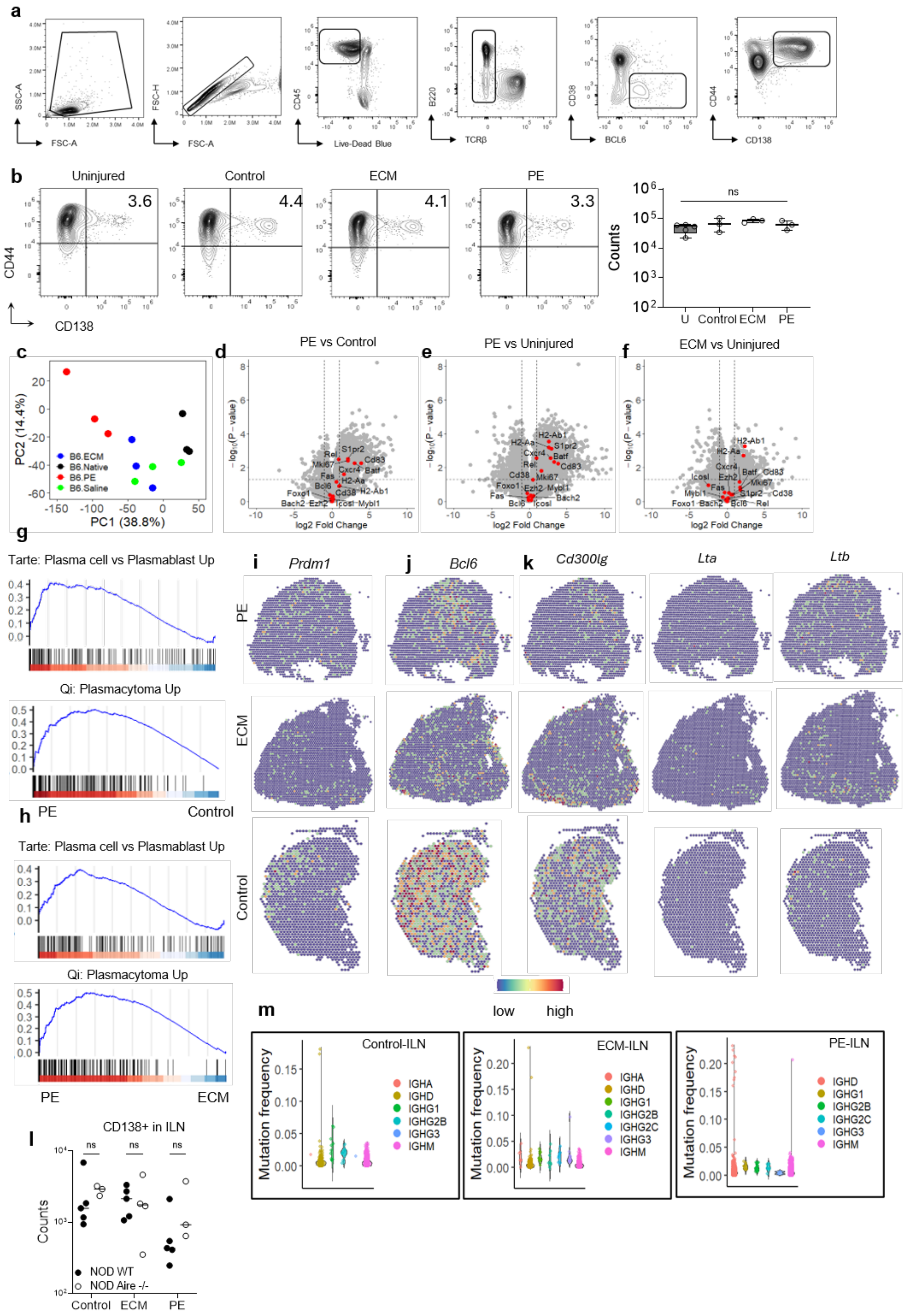

a
SSC-A
FSC-A
FSC-H
FSC-A
CD45
Live-Dead Blue
B220
TCRβ
CD38
BCL6
CD44
CD138
b
Uninjured
Control
ECM
PE
3.6
4.4
4.1
3.3
CD44
CD138
Counts
ns
U
Control
ECM
PE
c
PC2 (14.4%)
PC1 (38.8%)
B6.ECM
B6.Native
B6.PE
B6.Saline
d
PE vs Control
e
PE vs Uninjured
f
ECM vs Uninjured
log2 Fold Change
g
Tarte: Plasma cell vs Plasmablast Up
Qi: Plasmacytoma Up
h
PE
Control
Tarte: Plasma cell vs Plasmablast Up
Qi: Plasmacytoma Up
PE
ECM
i
Prdm1
j
Bcl6
k
Cd300lg
Lta
Ltb
PE
ECM
Control
low
high
l
CD138+ in ILN
ns
ns
ns
Counts
NOD WT
NOD Aire -/-
Control
ECM
PE
m
Control-ILN
ECM-ILN
PE-ILN
Mutation frequency
IGHA
IGHD
IGHG1
IGHG2B
IGHG2C
IGHG3
IGHM

**Supplemental Fig.6 | Ectopic germinal center formation after muscle trauma gives rise to plasma cells in tissues with microplastics**. (**a**) Gating strategy to identify CD138+ CD44+ plasma cells. (**b**) Splenic plasma cell counts in indicated treatments after volumetric muscle loss. (**c**) PCA plot of muscle tissue bulk RNA sequencing from uninjured mice and mice which received VML and treatments at 21dpi. (**d**) Differential expression of germinal center and plasma cell gene signature in tissue with microplastics compared to control injury and (**e**) uninjured tissue. (**f**) Notable absence of plasma cell markers in tissue with $ECM_{tx}$ as compared to uninjured muscle tissue. (**g**) Gene set enrichment analysis in tissue with microplastics compared to control injury and (**h**) $ECM_{tx}$ showing enrichment of genes relevant to plasma cell differentiation. (**i**) Spatial distribution of the germinal center B cell markers *Prdm1* and (**j**) *Bcl6* alongside the HEV marker *Cd300lg,* and TLS markers *Lta* and *Ltb* (**k**) in muscle tissue with indicated treatments. (**l**) Enumeration of plasma cells generated in inguinal lymph nodes of NOD WT and NOD *Aire*$^{-/-}$ mice 21 dpi. (**m**) Somatic hypermutation within plasma cells of indicated isotypes in inguinal lymph nodes of WT C57BL6 mice 21 dpi as analyzed by BCR sequencing.

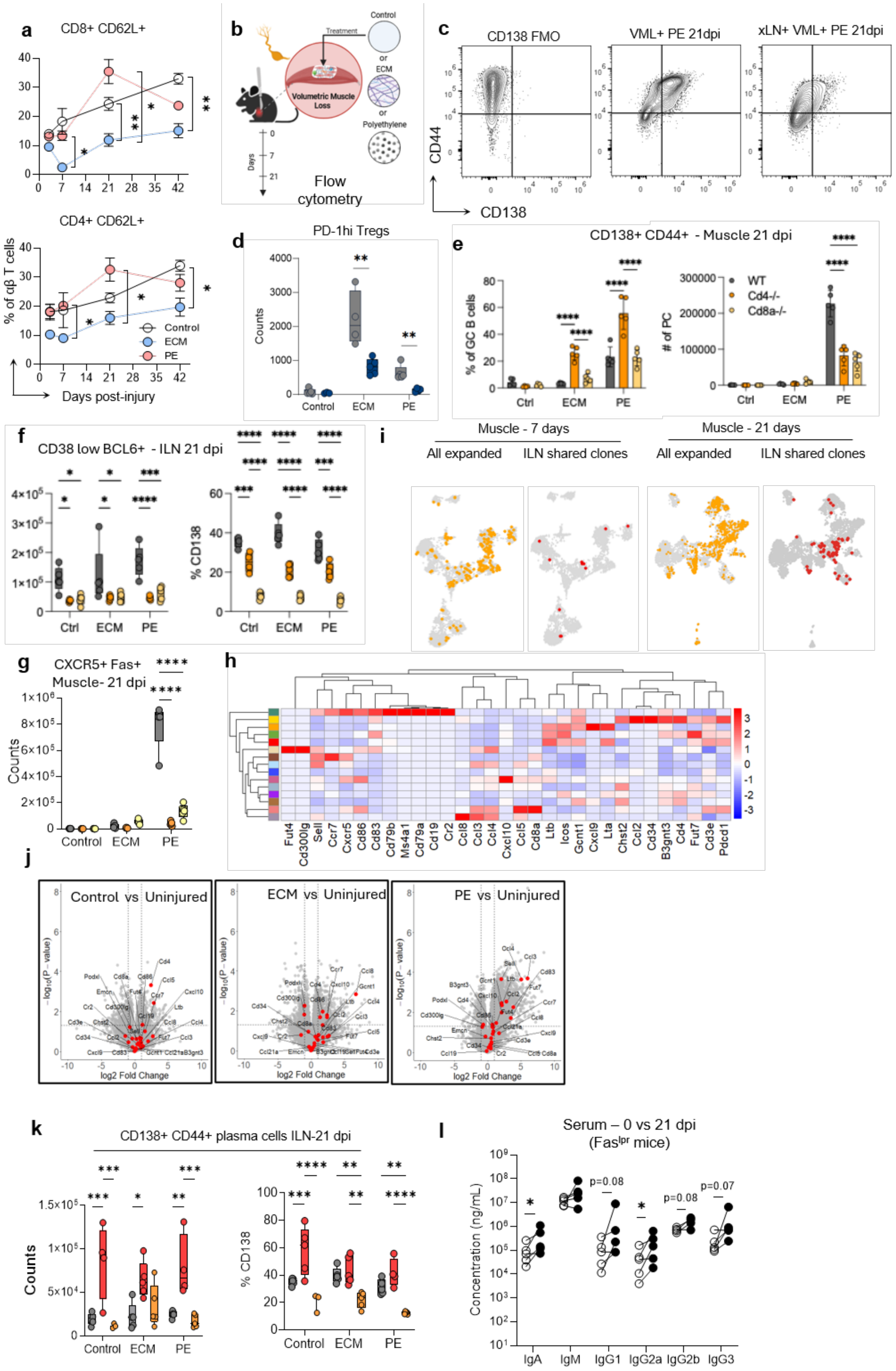

a
CD8+ CD62L+
CD4+ CD62L+
% of αβ T cells
Days post-injury
Control
ECM
PE
b
Treatment
Control
or
ECM
or
Polyethylene
Volumetric Muscle Loss
Days
Flow cytometry
c
CD138 FMO
VML+ PE 21dpi
xLN+ VML+ PE 21dpi
CD44
CD138
d
PD-1hi Tregs
Counts
Control
ECM
PE
e
CD138+ CD44+ - Muscle 21 dpi
% of GC B cells
# of PC
WT
Cd4-/-
Cd8a-/-
Ctrl
ECM
PE
f
CD38 low BCL6+ - ILN 21 dpi
% CD138
Ctrl
ECM
PE
i
Muscle - 7 days
Muscle - 21 days
All expanded
ILN shared clones
All expanded
ILN shared clones
g
CXCR5+ Fas+
Muscle- 21 dpi
Counts
Control
ECM
PE
h
Fut4
Cd300lg
Sell
Ccr7
Cxcr5
Cd86
Cd83
Cd79b
Ms4a1
Cd79a
Cd19
Cr2
Ccl8
Ccl3
Ccl4
Cxcl10
Ccl5
Cd8a
Ltb
Icos
Gcnt1
Cxcl9
Lta
Chst2
Ccl2
Cd34
B3gnt3
Cd4
Fut7
Cd3e
Pdcd1
j
Control vs Uninjured
ECM vs Uninjured
PE vs Uninjured
-log10(P-value)
log2 Fold Change
k
CD138+ CD44+ plasma cells ILN-21 dpi
Counts
% CD138
Control
ECM
PE
l
Serum – 0 vs 21 dpi
(Fas^lpr mice)
Concentration (ng/mL)
p=0.08
p=0.08
p=0.07
IgA
IgM
IgG1
IgG2a
IgG2b
IgG3

**Supplemental Fig.7 | Draining lymph node T cell dependence of plasma cell differentiation and regulation in damaged tissue.** (**a**) Retention of naïve like CD62L+ T cells in damaged tissue with indicated treatments over a time period of 42 days after VML. (**b**) Study design of experiments comparing plasma cell differentiation within injured tissue of mice with intact lymph nodes and those which underwent bilateral lymphadenectomy. (**c**) Representative flow cytometry contour plots showing plasma cells in injured tissue (PE group) of mice which underwent lymphadenectomy compared to mice with intact lymph nodes. (**d**) Enumeration of CD4+ PD-1+ Foxp3- T cells in damaged tissue at 21 dpi. (**e**) Fraction and counts of plasma cells in *Cd8a*$^{-/-}$ and *Cd4*$^{-/-}$ mice within injured tissue at 21 dpi. (**f**) Enumeration of TCRβ- CD38 low BCL6+ cells and fractions of CD138+ plasma cells in draining lymph nodes of WT, *Cd4*$^{-/-}$ and *Cd8a*$^{-/-}$ mice. (**g**) Enumeration of CD38 low BCL6+ GC-like B cells in muscle tissue in WT, *Cd4-/-* and *Cd8a-/-* mice at 21dpi. (**h**) Expression of putative markers for maintenance and support of tertiary lymphoid structures in T cell clusters in figure 3 at 21 dpi. (**i**) Clonal tracking of T cells between tissue and draining lymph nodes at 7 and 21 dpi using paired receptor sequencing. (**j**) Volcano plots showing TLS marker upregulation after VML at 21dpi in indicated groups compared to uninjured tissue by whole tissue bulk RNA sequencing. (**k**) Counts and abundance of CD138+ CD44+ plasma cells at 21dpi in draining lymph nodes of WT (grey), Fas$^{lpr}$ (red) and *Cd8a-/-* (orange) mice which received VML and indicated treatments. (**l**) Serum levels of indicated antibody isotypes at 21 dpi in Fas$^{lpr}$ mice which received PE.

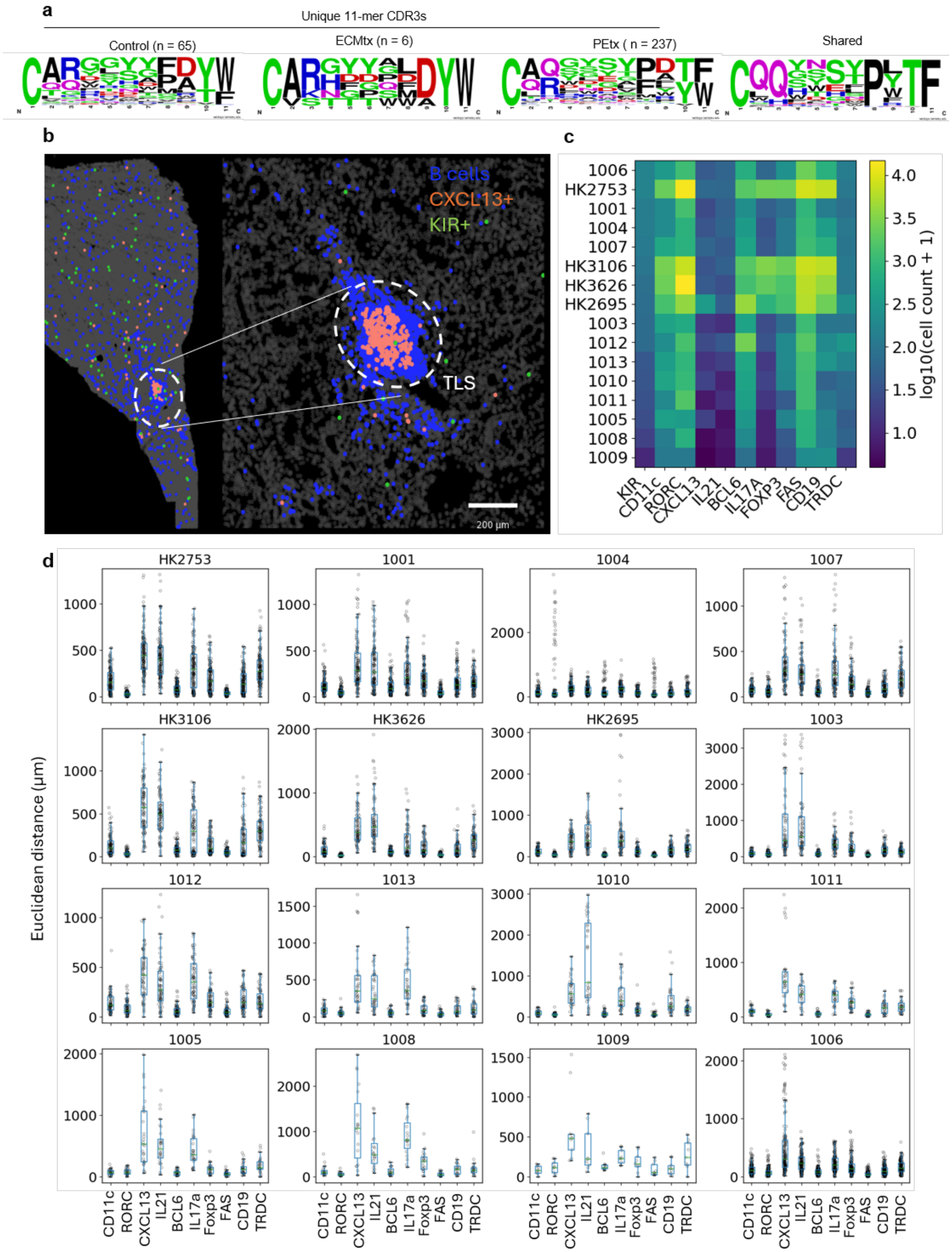


**Supplemental Fig.8 | B cell self-reactivity after VML and human KIR+ T cell recruitment to chronically inflamed damaged tissue** (**a**) Shared and unique B cell CDR3 sequences at 7dpi in indicated treatment groups. (**b**) Spatial distribution of KIR+ cells around TLS in a human with diabetic kidney disease and glomerular damage (**c)** Enumeration of DC, T cell and B cell markers across 15 spatial transcriptomic sections obtained from dataset published by Dumoulin et.al[45] (**d)** Nearest neighbor analysis showing physical proximity of KIR expressing T cells to dendritic cells and *RORC* expressing αβ T cells.
